# Dopamine and norepinephrine differentially mediate the exploration-exploitation tradeoff

**DOI:** 10.1101/2023.01.09.523322

**Authors:** Cathy S. Chen, Dana Mueller, Evan Knep, R. Becket Ebitz, Nicola M. Grissom

## Abstract

The catecholamines dopamine (DA) and norepinephrine (NE) have been implicated in neuropsychiatric vulnerability, in part via their roles in mediating the decision making processes. Although the two neuromodulators share a synthesis pathway and are co-activated, they engage in distinct circuits and roles in modulating neural activity across the brain. However, in the computational neuroscience literature, they have been assigned similar roles in modulating the exploration-exploitation tradeoff. Revealing how each neuromodulator contributes to this explore-exploit process is important in guiding mechanistic hypotheses emerging from computational psychiatric approaches. To understand the differences and overlaps of the roles of dopamine and norepinephrine in mediating exploration, a direct comparison using the same dynamic decision making task is needed. Here, we ran mice in a restless bandit task, which encourages both exploration and exploitation. We systemically administered a nonselective DA antagonist (flupenthixol), a nonselective DA agonist (apomorphine), a NE beta-receptor antagonist (propranolol), and a NE beta-receptor agonist (isoproterenol), and examined changes in exploration within subjects across sessions. We found a bidirectional modulatory effect of dopamine receptor activity on exploration - increasing dopamine activity decreased exploration and decreasing dopamine activity increased exploration. The modulation of exploration via beta-noradrenergic activity was mediated by sex. Computational model parameters revealed that dopamine modulation affected exploration via decision noise and norepinephrine modulation via outcome sensitivity. Together, these findings suggested that the mechanisms that govern the transition between exploration and exploitation are sensitive to changes in both catecholamine functions and revealed differential roles for NE and DA in mediating exploration.

**Significance Statement:** Both dopamine (DA) and norepinephrine (NE) has been implicated in the decision making process. Although these two catecholamines have shared aspects of their biosynthetic pathways and projection targets, they are thought to exert many core functions via distinct neural targets and receptor subtypes. However, the computational neuroscience literature often ascribes similar roles to these catecholamines, despite the above evidence. Resolving this discrepancy is important in guiding mechanistic hypotheses emerging from computational psychiatric approaches. This study examines the role of dopamine and norepinephrine on the explore-exploit tradeoff. By testing mice, we were able to compare multiple pharmacological agents within subjects, and examine source of individual differences, allowing direct comparison between the effects of these two catecholamines in modulating decision making.

## Introduction

Dysfunctions in cognitive processes, particularly within the domain of executive function, are implicated in numerous neuropsychiatric disorders (Elliott, 2003; Malloy-Diniz et al., 2017; Grissom and Reyes, 2019). One essential aspect of executive function is value-based decision making. Decision making involving exploring and learning in an uncertain environment has been a focus in computational neuroscience, which has developed approaches for discovering latent structures in decision making strategies (Rothenhoefer et al., 2017; Ebitz et al., 2018; Chen et al., 2021a, 2021b). Many computational models for value-based decision making distinguish two essential latent processes: exploration and exploitation (Sutton and Barto, 1998; Niv et al., 2002; Daw et al., 2006; Jepma et al., 2020; Wilson et al., 2021). Exploration is the ubiquitous learning process across species that allows the discovery of the best action to take in an uncertain environment (Ebitz et al., 2018; Gershman, 2019; Chen et al., 2021b; Wilson et al., 2021). Once a favorable action or option is discovered, exploitation, i.e. repeating a rewarding action, is necessary to obtain the best rates of reward. However, exploitation must be balanced with continued exploration as environments and their reward probabilities change. Dysregulation in this exploration-exploitation tradeoff is implicated in the phenotypes of numerous neuropsychiatric disorders and challenges, including schizophrenia, autism spectrum disorders, addictions, and chronic stress (Kim et al., 2007; Mussey et al., 2015; Morris et al., 2016; Addicott et al., 2020; Kaske et al., 2022). Understanding the fundamental mechanisms impacting the computations in our brain that underlie the balance between exploration and exploitation could help identify critical circuits that are associated with differential risk factors for neuropsychiatric disorders and open avenues for novel interventions for executive function challenges.

Many neuropsychiatric challenges are associated with dysfunction in the catecholamines dopamine and norepinephrine (Kobayashi, 2001; Ressler and Nemeroff, 2001; Aston-Jones and Cohen, 2005; Bouret and Sara, 2005; Money and Stanwood, 2013; Addicott et al., 2020; Williams et al., 2021). These neuromodulators are well-positioned to carry information about the state of the environment and influence behavior outputs via downstream control of action selection (Cohen et al., 2007; Wilson et al., 2021). Indeed, dopamine and norepinephrine have each separately been implicated in mediating the exploration-exploitation tradeoff (Seamans and Yang, 2004; Aston-Jones and Cohen, 2005; Bouret and Sara, 2005; Cohen et al., 2007; Redish et al., 2007; Wilson et al., 2021; Cremer et al., 2022). While dopamine and norepinephrine share biosynthetic pathways and are co-activated under states of arousal, they act through partially separable circuits and have distinct neural targets (Ranjbar-Slamloo and Fazlali, 2019). However, in the computational neuroscience literature, dopamine and norepinephrine have largely been ascribed similar or even identical roles in mediating the explore-exploit tradeoff despite of the distinct pharmacological profiles.

We have previously shown that multiple latent cognitive processes, such as the reinforcement learning model-derived parameters of learning rate and decision noise, can describe differences in the level of exploration between groups (Chen et al., 2021b). Dopamine has often been implicated in mediating reward prediction errors and value-based learning (Schultz, 1998, 2013; Daw et al., 2006; Niv, 2009) while norepinephrine in modulating arousal, attention, and value assessment (Aston-Jones and Cohen, 2005; Yu and Dayan, 2005; Amemiya et al., 2016), but more recently both neuromodulators have been implicated in the process of action selection (Humphries et al., 2012; Warren et al., 2017). Therefore, it is unknown whether dopamine and norepinephrine govern this exploration-exploitation tradeoff via distinct or similar latent cognitive processes because they have been largely studied in this context in isolation (Cremer et al., 2022).

To uncover the distinct roles of dopamine and norepinephrine in the exploration-exploitation tradeoff, we conducted within-subjects pharmacological manipulations for these two neuromodulators to examine their effect on exploration and the latent cognitive parameters that influence exploration, in the same restless bandit task. We pharmacologically up- and down-regulated dopaminergic and noradrenergic receptor activity and compared the modulatory effects on exploration in the same dynamic decision making task, a spatial restless two-armed bandit (Ebitz et al., 2018; Chen et al., 2021b). We found a bidirectional modulatory effect of dopamine receptor activity on the level of exploration. Increasing dopamine activity decreased exploration and decreasing dopamine activity increased exploration. Beta-noradrenergic receptor activity also modulated exploration but the modulatory effect was mediated by sex. Contrary to dopamine modulating choice solely by changing value-based learning for outcomes, via reinforcement learning modeling we find that dopamine mediates exploration by changing decision noise, i.e. the precision of value-based choice selection. In contrast, noradrenergic activity surprisingly influenced exploration by changing value-based learning for outcomes in a sex dependent manner. These data suggest differential roles of dopamine and norepinephrine in mediating exploration, and complex roles for both catecholamines in signaling reward.

## Materials and Methods

### Animals

Thirty-two BL6129SF1/J mice (16 males and 16 females) were obtained from Jackson Laboratories (stock #101043). Mice arrived at the lab at 7 weeks of age and adapted to 0900-2100 hours reversed light cycle throughout the testing phase. Mice were housed in groups of four with ad libitum access to water while being mildly food restricted (85-95% of free feeding weight) before the start of the experiment (12 weeks) and during the experiment. All animals were cared for according to the guidelines of the National Institution of Health and experimental protocols were approved by the Institutional Animal Care and Use Committee (IACUC) of the University of Minnesota.

### Apparatus

Sixteen identical triangular touchscreen operant chambers (Lafayette Instrument Co., Lafayette, IN) were used for training and testing. Two walls were black acrylic plastic. The third wall housed the touchscreen and was positioned directly opposite the magazine. The magazine provided liquid reinforcers (50% Ensure) delivered by a peristaltic pump, typically 7ul (280 ms pump duration). ABET-II software (Lafayette Instrument Co., Lafayette, IN) was used to program operant schedules and to analyze all data from training and testing.

### Behavioral task

#### Two-armed spatial restless bandit task

Animals were trained to perform a two-armed spatial restless bandit task in the touchscreen operant chamber. Each trial, animals were presented with two identical squares on the left and right side of the screen. Nose poke to one of the target locations on the touchscreen was required to register a response. Each location is associated with some probability of reward, which changes independently over time. For every trial, there is a 10% chance that the reward probability of a given arm will increase or decrease by 10%. All the walks were generated randomly with a few criteria: 1) the overall reward probabilities of two arms are within 20% of each other, preventing one arm being overly better than the other, 2) the reward probability cannot go down to 0% or go up to 100%, 3) there are no 30 consecutive trials where the reward probabilities of both arms are lower than 20% to ensure motivation.

Animals ran for 6 control sessions and 6 drug sessions for each pharmacological manipulation. The wash-out period between each drug condition is 3 days. Each session consisted of 300 trials. Within each session, animals completed either 300 trials or spent a maximum of two hours in the operant chamber. On average across all sessions, animals performed 276.5 trials with a standard deviation of 8.6 trials (male average: 253.7 trials, std = 15.4; female average 299.3 trials, std = 0.74. Data were recorded by the ABET II system and was exported for further analysis. All computational modeling was conducted using python.

### Drug administration

To assess the effect of norepinephrine and dopamine receptor activity on explore behaviors, we used the following four different drugs to increase or decrease beta-noradrenergic or dopaminergic receptor activity systemically: a beta-noradrenergic receptor agonist isoproterenol (0.3mg/kg), a beta-noradrenergic receptor antagonist propranolol (5mg/kg), a non-selective dopamine receptor agonist apomorphine (0.1mg/kg), and a non-selective dopamine receptor antagonist flupenthixol (0.03mg/kg).

All drugs were fully dissolved in 0.9% saline and protected from light. Animals received intraperitoneal (IP) injection of drug or saline (control) at an injection volume of 5ml/kg right before the experiment with the exception of apomorphine being administered 20 minutes before the experiment due to the immediate drug influence of locomotor behaviors. Drug and saline were administered in alternating sessions using a within-subject so that every animal received each of the four drugs. The dopamine receptor agonist and antagonist were administered counterbalanced across animals. Half of the animals (n = 8) were randomly selected to receive apomorphine first, the other received flupenthixol first. Since this experiment prioritized using a within-subject design to understand the effect of each drug within individual animals and the source of individual differences, the length of study only limited us to test one dose for each of the four drugs. Therefore, dosage chosen was based on previous studies on the effect of drugs on cognitive functions (Fernandez-Tome et al., 1979; Cabib et al., 1984; Goldschmidt et al., 1984; Ichihara et al., 1988; Nakamura et al., 1998; Grigoryan, 2012; Cinotti et al., 2019). Doses for each drug were chosen from previous studies as the lowest doses necessary to produce alterations in cognitive functions including decision making, learning and exploration with minimal influence on locomotion.

### Data Analysis

#### General analysis techniques

Data were analyzed with custom PYTHON, and Prism 8 scripts. Generalized linear mixed models (GLMMs), ANOVA, and t-test were used to determine the fixed effects of drug, sex, and interaction term, accounting for random effects of animal identity and session, unless otherwise specified. P values were compared against the standard ɑ = 0.05 threshold. The sample size is n = 32 (16 males and 16 females) for all statistical tests. No animal was excluded from the experiment. All statistical tests used and statistical details were reported in the results. All figures depict mean ± SEM.

#### Generalized Linear Mixed Models (GLMMs)

In order to determine whether drug and sex predicted the reward acquisition performance, response time, probability of win-stay, probability of lose-shift, outcome sensitivity, probability of exploration, we fitted a series of generalized linear mixed model to examine the fixed effects of drug, sex, and interaction term, with animal identity and sessions as random effects.

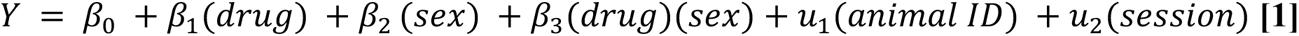

where Y is the dependent variable (obtained reward, response time, win-stay, lose-shift, outcome sensitivity, exploration). In this model, ꞵ1 describes the main effect of drug and ꞵ2 describes the main effect of sex. ꞵ3 captures any interaction effect between drug and sex. u1 and u2 capture the random effect of animal identity and session respectively.

#### Outcome Sensitivity

In order to examine how much of the switching behavior was sensitive to reward outcome, we calculated the difference between switching given no reward and switching given reward control by the total amount of switching. If switching behavior was not affected by the outcome at all, the outcome sensitivity should be close to zero.

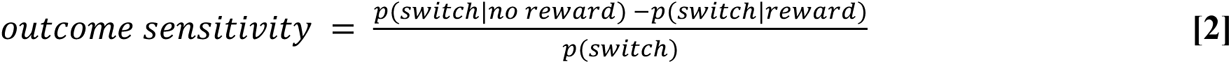

### Hidden Markov model (HMM)

In order to identify when animals were exploring, we fit a Hidden Markov model (HMM) to the animals’ choice sequence. Our model consisted of two hidden states - explore state and exploit state, which are defined by the probability of making each choice (k, out of N_k_ possible options) and the probability of transitioning from one state to another. When an HMM model is fitted to animals’ choices, it estimates a transition matrix, which is the mapping between every state at one time point and every state at some future point of time. The HMM model fitting is the process of estimating how the distribution over states will evolve over time. The transition matrix estimates the probability of each type of transition between states. The HMM models exploration as a uniform distribution over choices, i.e. the emissions model for the explore state was uniform across options. This is the maximum entropy distribution of a categorical variable, which makes the few assumptions of true distribution of choices during exploration and therefore does not bias the model towards or away from any particular type of high-entropy choice period. In the exploitation state, subjects repeatedly sample the same choice, therefore the exploit states only permit one choice, i.e. exploit-left state only emits left choices and exploit-right state only emits right choice.

The latent states in this model are Markovian, meaning that they are time-independent. They depend only on the most recent state. The HMM model estimates a transition matrix to fit the behavior by mapping of past and future states. This matrix is a system of stochastic equations describing the 1-time-step probability of transitioning between every combination of states. In our model, there were 3 possible states (2 exploit states and 1 explore state). The parameters were tied across exploit states such that each exploit state had the same probability of beginning (from exploring) and of sustaining itself. Transitions out of the exploration, into exploitative states, were also tied. The model also assumed that the mice had to pass through exploration in order to start exploiting a new option, even if only for a single trial. Through fixing the emissions model, constraining the structure of the transmission matrix, and tying the parameters, the final HMM had only two free parameters: one corresponding to the probability of exploring, given exploration on the last trial, and one corresponding to the probability of exploiting, given exploitation on the last trial.

The model was fit via expectation-maximization using the Baum Welch algorithm (Bilmes, 1998). This algorithm finds a (possibly local) maxima of the complete-data likelihood. A complete set of parameters θ includes the emission and transition models, discussed already, but also initial distribution over states, typically denoted as PI. Because the mice had no knowledge of the environment at the first trial of the session, the initial distribution started with p (explore state) = 1. The algorithm was reinitialized with random seeds 10 times, and the model that maximized the observed (incomplete) data log likelihood across all the sessions for each animal was ultimately taken as the best. To decode latent states from choices, we used the Viterbi algorithm to discover the most probable posteriori sequence of latent states.

#### Reinforcement learning (RL) models

We fitted five reinforcement learning (RL) models that could potentially characterize animals’ choice behaviors, with the structure of each RL model detailed below. AIC weights were calculated from AIC values across all treatment groups and compared across models to determine the best model with the highest relative likelihood. Model agreement was calculated as the probability of choice given model to demonstrate how well the best fit model captures actual choices.

The first model (random) assumes that animals choose between two arms randomly with some overall bias for one side over the other. This choice bias for choosing left side over right side is captured with a parameter *b.* The probability of choosing left side on trial *t* is:

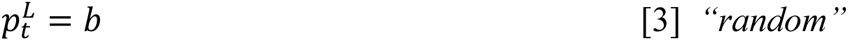

The second model is a noisy win-stay lose-shift (WSLS) model that adapts choices with regards to outcomes. This model assumes a win-stay lose-shift policy that is to repeat a rewarded choice and to switch to the other choice if not rewarded. Furthermore, this model includes a parameter ɛ that captures the level of randomness, allowing a stochastic application of the win-stay lose-shift policy. The probability of choosing arm *k* on trial *t* is:

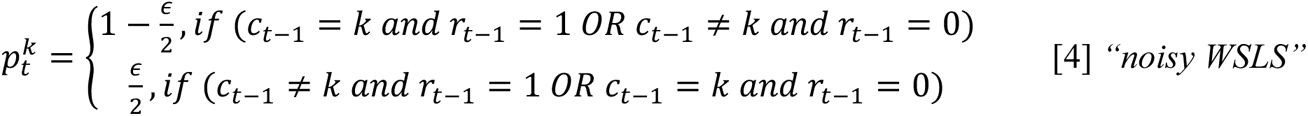

c_t_ indicates the choice on trial *t* and r_t_ is a binary variable that indicates whether or not trial *t* was rewarded.

The third model (RL) is a basic delta-rule reinforcement learning (RL) model. This two-parameter model assumes that animals learn by consistently updating Q values, which are values defined for options (left and right side). These Q values, in turn, dictate what choice to make next. For example, in a multi-armed bandit task, 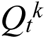 is the value estimation of how good arm *k* at trial *t*, and is updated based on the reward outcome of each trial:

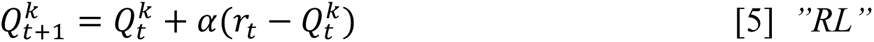

In each trial, 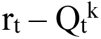 captures the reward prediction error (RPE), which is the diffe-rence between expected outcome and the actual outcome. The parameter *a* is the learning rate, which determines the rate of updating RPE. With Q values defined for each arm, choice selection on each trial was performed based on a Softmax probability distribution:

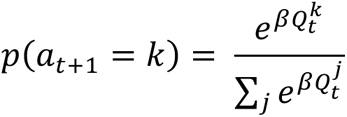

where inverse temperature β determines the level of random decision noise.

The fourth model (*RLCK*) incorporates a choice updating rules in addition to the value updating rule in model 3. The model assumes that choice kernel, which captures the outcome-independent tendency to repeat a previous choice, also influences decision making. The choice kernel updating rule is similar to the value-updating rule:

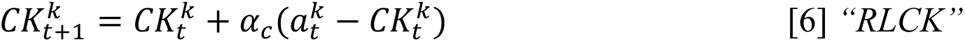

where a_t_^k^ is a binary variable that indicates whether or not arm *k* was chosen on trial *t* and *a*_t_ is choice kernel updating rate, characterizing choice persistence. The value and choice kernel term were combined to compute the probability of choosing arm *k* on trial *t*:

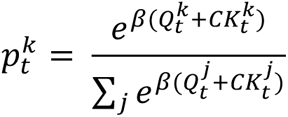

where β is the inverse temperature, capturing the decision noise outside value and choice bias.

The fifth model (*RLγ)* is an asymmetrical learning model that incorporates an asymmetric learning scalar parameter that scales the learning rate on the trials where there is no reward. The choice selection is also based on a Softmax probability distribution where the inverse temperature determines the decision noise in the system.

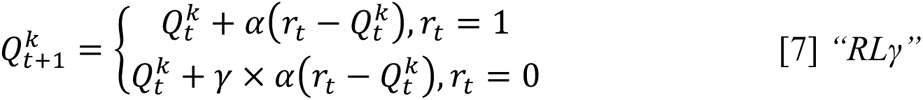

The sixth model (*RLCKγ*) is the combination of the fourth and the fifth model, incorporating both asymmetrical learning and choice kernel. This model included a learning rate, an asymmetric learning scalar, a choice kernel, and an inverse temperature parameter.

## Results

Aged-matched adult male and female wildtype mice (n = 32, 16 males and 16 females, strain B6129SF1/J) were trained to perform a restless two-armed spatial bandit task in touch-screen operant chambers (**Figure 1A**). In this task, animals were presented with two identical visual targets (squares) on the left and right side of the screen during each trial. They indicated their choices by nose poking at one of the two target locations, each of which provided some probability of reward that changed independently and randomly over time (**Figure 1B)**. The dynamic reward contingency of this task naturally encourages the animal to balance between exploiting a favorable option when it is found and exploring to gain information about potential better alternatives. This task has been employed in rodents and primates as well as human subjects to understand learning and exploration and have successfully revealed divergent exploration strategies (Ebitz et al., 2018; Grossman et al., 2020; Chen et al., 2021b). In this study, we adopted this task to understand the modulatory effect of catecholamine receptor activity on the transition between exploration and exploitation.

**Figure 1.**
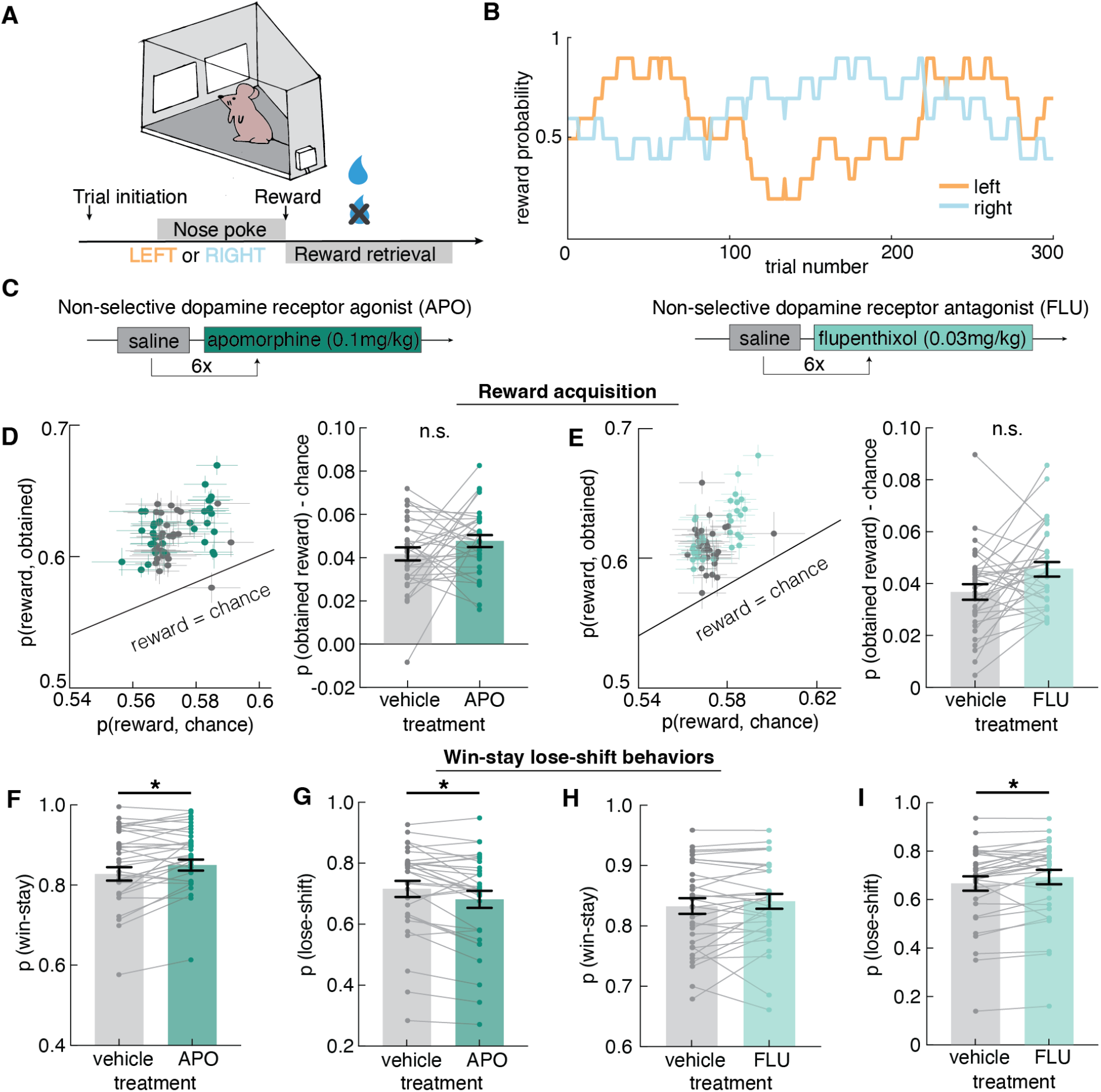
Modulation of dopamine receptor activity affected reward-directed behaviors. Up-regulating dopamine activity increased stickiness in choice behaviors regardless of outcomes. **A)** Schematic of the mouse touchscreen chamber and the trial structure of the two-armed spatial restless bandit task. **B)** An example of the dynamic reward contingency showing the changing reward probabilities associated with each option over 300 trials. **C)** The dopaminergic drug administration schedule and drug dosage used to modulate dopamine receptor activity. A non-selective dopamine receptor agonist apomorphine (0.1mg/kg) and a non-selective dopamine receptor antagonist flupenthixol (0.03mg/kg) were systemically administered to respectively up- and down-regulate dopamine receptor activity. 0.9% saline was used as the vehicle control. Control and drug sessions were interleaved and repeated for six sessions each. **D)** (left) Average probability of obtaining reward compared to the chance level probability of reward for each animal under apomorphine (APO) and vehicle (dot). Error bars depict mean ± SEM across sessions for each animal. (right) probability of obtaining reward over chance level across vehicle and APO condition. **E)** (left) Average probability of obtaining reward compared to the chance level probability of reward for each animal under flupenthixol (FLU) and vehicle (dot). Error bars depict mean ± SEM across sessions for each animal. (right) probability of obtaining reward over chance level across vehicle and FLU condition. **F)** Average probability of win-stay on vehicle and APO condition. **G)** Average lose-shift on vehicle and APO condition. **H)** Average probability of win-stay on vehicle and FLU condition. **I)** Average lose-shift on vehicle and FLU condition. * indicates p < 0.05. Graphs depict mean ± SEM across animals unless specified otherwise.

To examine the distinct and/or overlapping roles of dopamine and norepinephrine in modulating exploration, we tested the systemic effect of a non-selective dopamine receptor agonist apomorphine (0.1mg/kg), a non-selective dopamine receptor antagonist flupenthixol (0.03mg/kg), a beta-noradrenergic receptor agonist isoproterenol (0.3mg/kg), and a beta-noradrenergic receptor antagonist propranolol (5mg/kg) on exploration. The experiment used a within-subject design, where all animals received each of the four drugs tested. This design allowed us to directly compare the effect of each drug within individual animals and dissociate drug effects from potential cohort differences. Since each animal received all four drugs, the length of study did not allow us to test multiple doses for each drug. Therefore, we selected the dosages of the drugs based on numerous previous studies on the role of dopamine and norepinephrine in cognitive processes (Fernandez-Tome et al., 1979; Cabib et al., 1984; Goldschmidt et al., 1984; Ichihara et al., 1988; Nakamura et al., 1998; Grigoryan, 2012; Cinotti et al., 2019). Animals were intraperitoneally (IP) administered either saline (control) or treatment with 5 mg/kg of propranolol, 0.3mg/kg of isoproterenol, or 0.03 mg/kg of flupenthixol immediately prior to the bandit task or 0.1mg/kg of apomorphine 30 minutes prior to the bandit task. Saline and drug treatments were administered interleaved by session (**Figure 1C, 2A**). Mice performed 12 sessions under each pharmacological condition (6 control and 6 drug treatment sessions). Each drug effect was compared to its own vehicle control collected in the interleaved sessions.

To examine whether mice had learned to perform the task and were able to sustain reward, we first calculated the average probability of obtaining reward across sessions in all control and drug groups. It’s worth noting that in this task, there is no asymptotic performance because the animals need to constantly adapt to the changing reward contingency and any one option is not always the best choice to make. At any given time, an animal could decide to repeat a favorable option or to explore the options. The performance is best measured by the amount of reward acquired. Because reward schedules were dynamic and stochastic, sessions could differ slightly in the amount of reward that was available. We therefore compared the probability of actual obtained reward against the probability of reward if chosen randomly (chance). In the dopamine modulation condition, both control groups (saline) and treatment groups (apomorphine and flupenthixol) were able to obtain reward more frequently than chance (**Figure 1D, E**, one-sample t test, vehicle control (apomorphine): t(31) = 13.96, p<0.0001; apomorphine: t(31)= 17.24, p<0.0001; vehicle control (flupenthixol): t(31)=12.36, p<0.0001; flupenthixol: t(31)=16.48, p<0.0001). When modulating beta-noradrenergic receptor activity, both control groups (saline) and treatment groups (isoproterenol and propranolol) were able to obtain reward more frequently than chance (**Figure 2B, D**, one-sample t test, vehicle control (isoproterenol): t(31)=10.01, p<0.0001; isoproterenol: t(31)=13.84, p<0.0001; vehicle control (propranolol): t(31)=10.60, p<0.0001; propranolol: t(31)=20.08, p<0.0001). These results suggested that animals were not choosing randomly. The animals demonstrated a strong understanding of the task and were able to effectively obtain rewards above chance under baseline (vehicle control) and all drug treatments.

**Figure 2.**
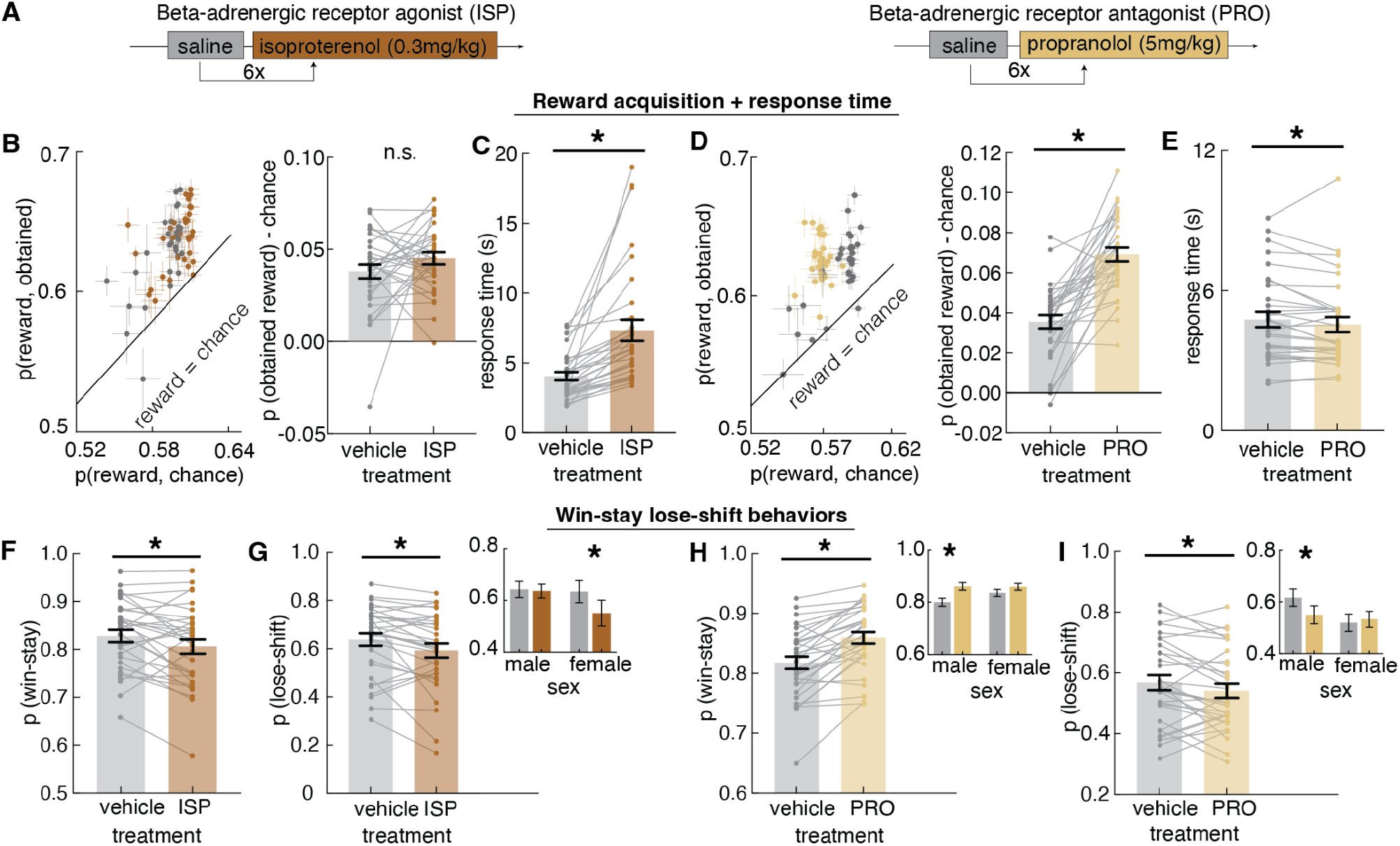
The modulatory effects of beta-noradrenergic receptor activity on reward-directed behavior was also influenced by sex. **A)** The noradrenergic drug administration schedule and drug dosage used to modulate noradrenergic receptor activity. A beta-noradrenergic receptor agonist isoproterenol (0.3mg/kg) and a beta-noradrenergic receptor antagonist propranolol (5mg/kg) were systemically administered to respectively up- and down-regulate norepinephrine activity. 0.9% saline was used as the vehicle control. Control and drug sessions were interleaved and repeated for six sessions each. **B)** (left) Average probability of obtaining reward compared to the chance level probability of reward for each animal under isoproterenol (ISP) and vehicle (dot). Error bars depict mean ± SEM across sessions for each animal. (right) probability of obtaining reward over chance level across vehicle and ISP. **C)** Average response time under ISP and vehicle. ISP significantly increased response time. **D)** (left) Average probability of obtaining reward compared to the chance level probability of reward for each animal under vehicle and propranolol (PRO). Error bars depict mean ± SEM across sessions for each animal. (right) probability of obtaining reward over chance level across PRO (light brown) and vehicle (gray). Average response time under PRO and vehicle. PRO significantly decreased response time. **E)** Average probability of win-stay under ISP and vehicle. **G)** Average probability of lose-shift under ISP and vehicle. (Inset) probability of lose-shift across treatments across sexes. Decrease in lose-shift under ISP was primarily driven by changes in females (interaction term). **H)** Average probability of win-stay under PRO and vehicle. (Inset) probability of win-stay across treatments across sexes. Increase in win-stay under PRO was primarily driven by changes in males (interaction term). **I)** Average probability of lose-shift under PRO and vehicle. (inset) probability of lose-shift across treatments across sexes. Decrease in lose-shift under PRO was primarily driven by changes in males (interaction term). * indicates p < 0.05. Graphs depict mean ± SEM across animals unless specified otherwise.

Next, we asked whether manipulating dopaminergic receptor activity influenced their reward acquisition performance. We used a generalized linear mixed model (GLMM) to estimate effects and interactions of drug treatments and sex, with session and animal identity as random effects **(Equation 1**, see methods). Increasing dopamine receptor activity with apomorphine administration did not significantly alter their reward acquisition performance (**Figure 1D**, GLMM, main effect of drug, p= 0.11, ꞵ1 = −0.017). Antagonizing dopamine receptor activity with flupenthixol administration also did not significantly alter reward acquisition performance (**Figure 1E**, GLMM, main effect of drug, p= 0.28, ꞵ1 = −0.008). Increasing beta-noradrenergic receptor activity with isoproterenol administration did not affect the amount of reward obtained (**Figure 2B**, GLMM, main effect of drug, p= 0.93, ꞵ1 = −0.0005). However, when decreasing beta-noradrenergic receptor activity with propranolol administration, animals obtained significantly more reward (**Figure 2D**, GLMM, main effect of drug, p < 0.0001, ꞵ1 = −0.497). Even though manipulating dopamine receptor activity and increasing beta-noradrenergic receptor activity did not affect how much overall obtained reward, we considered the possibility that the tuning of catecholamine receptor activity affected latent exploration strategies underlying choice patterns, but not an observable change in the level of accuracy. Our previous study demonstrated that similar reward acquisition performance can be achieved via divergent explore strategies (Chen et al., 2021b).

Pharmacological manipulations of both dopaminergic and noradrenergic receptor activity have been shown to have a dose-dependent effect on motor functions (Goldschmidt et al., 1984; Weed and Gold, 1998). In this study, dosage selection was based on previous literature examining cognitive functions and a low dose was used to have influence on cognition but avoid motor function impairment (Nakamura et al., 1998; Cinotti et al., 2019). One commonly used behavioral metric to examine motor function in a cognitive task is response time. Therefore, we calculated the response time, which was the time elapsed between the onset of choices and nose poke response to make a choice, as recorded by the touchscreen operant chamber. If the administration of drugs impaired motor function, we would expect to see longer response time in the drug group compared to vehicle control. Upregulating dopamine receptor activity with apomorphine did not significantly influence the response time (GLMM, main effect of drug, p=0.77, ꞵ1 = 0.10). Similarly, downregulating dopamine receptor activity with flupenthixol did not significantly alter animals’ response time (GLMM, main effect of drug, p=0.96, ꞵ1 = −0.01). Modulating beta-noradrenergic receptor activity resulted in bidirectional changes in response time. When increasing beta-noradrenergic receptor activity with isoproterenol, both males and females took significantly longer to respond (**Figure 2C**, GLMM, main effect of drug, p<0.0001, ꞵ1 = −3.51). Decreasing beta-noradrenergic receptor activity with propranolol significantly decreased the response time (**Figure 2E**, GLMM, main effect of drug, p=0.04, ꞵ1 = 0.51), primarily driven by response time reduction in females under propranolol compared to vehicle controls (GLMM, interaction term, p<0.009, ꞵ3 = −0.57). Faster response time under propranolol could be associated with more reward obtained - propranolol was the only drug treatment that altered reward acquisition performance among the four drugs tested. Previous studies have linked response time with the complexity of the strategy used (Kool et al., 2010; Filipowicz et al., 2019; Chen et al., 2021b). Faster response time under propranolol might also point to the possibility that decreasing beta-noradrenergic receptor activity brought on changes in strategies that took shorter time to execute. Isoproterenol was a highly significant predictor for lower response time in both male and female mice, implicating the potential drug effect on motor behaviors.

### Opposing modulatory effect of dopamine and norepinephrine receptor activity on win-stay lose-shift behaviors

The role of dopamine has been heavily studied and shown to be a key contributor to value-based decision making and reinforcement learning (Seamans and Yang, 2004; Redish et al., 2007; Niv, 2009; Floresco, 2013). Although modulating dopamine receptor activity did not significantly influence the overall amount of reward obtained or the response time, it is possible that apomorphine and flupenthixol exerted influence on how animals adapted their choices to reward outcome. To understand how increasing and decreasing dopamine activity influenced animals’ reward-driven choice behaviors, we examined the probability of win-stay (repeating a choice when it was rewarded on the previous trial) and the probability of lose-shift (switching to the other choice when not rewarded). We found that when increasing dopamine receptor activity with apomorphine administration, both males and females had increased win-stay behavior (**Figure 1F**, GLMM, main effect of drug, p=0.011, ꞵ1= −0.02) and decreased lose-shift behavior (**Figure 1G**, GLMM, main effect of drug, p=0.016, ꞵ1= 0.034). This result suggested that apomorphine increased the overall “stickiness” or inflexibility of choice and more stay behaviors regardless of the outcome.

There was no significant change in the probability of win-stay when the animals were administered flupenthixol (**Figure 1H**, GLMM, main effect of drug, p=0.27, ꞵ1= −0.001). It’s worth noting that there was main effect of sex in the probability of win-stay, with females having a higher probability of win-stay, in both DA agonist and antagonist manipulations (GLMM, main effect of sex, apomorphine: p=0.065, ꞵ2 = −0.054; flupenthixol: p=0.01, ꞵ2 =-0.05). The results also revealed an opposing effect from dopamine agonist that antagonizing dopamine receptor activity with flupenthixol administration increased the probability of lose-shift (**Figure 1I**, GLMM, main effect of drug, p=0.04, ꞵ1 = −0.014).

To control for the total amount of switching, we calculated outcome sensitivity (**Equation 2**, see Methods) to understand how much of the switching behavior was outcome-sensitive. The result suggests that there were no significant changes in outcome sensitivity when animals were administered apomorphine (GLMM, main effect of drug, p=0.22, ꞵ1 = −0.08). However, administration of flupenthixol significantly increased outcome sensitivity across both sexes (**Figure 1-1A**, GLMM, main effect of drug, p=0.006, ꞵ1= −0.06) and there was more increase in outcome sensitivity in males (GLMM, interaction term, p=0.027, ꞵ3= −0.10).

Modulating beta-noradrenergic receptor activity also influenced win-stay and lose-shift behaviors. Increasing beta-noradrenergic activity with isoproterenol administration decreased the probability of win-stay compared to vehicle control (**Figure 2F**, GLMM, main effect of drug, 0.009, ꞵ1=0.007). Isoproterenol administration also decreased the probability of lose-shift in both sexes (**Figure 2G**, GLMM, main effect of drug, p=0.0007, ꞵ1=0.083), primarily driven by significant decrease in lose-shift in females under isoproterenol (**Figure 2G inset,** GLMM, interaction term, p=0.004, ꞵ3=-0.077). Outcome sensitivity analysis revealed a significant decrease in outcome sensitivity when administer isoproterenol compared to vehicle control (**Figure 1-1B**, GLMM, main effect of drug, p=0.021, ꞵ1 = 0.134). In contrast, decreasing beta-noradrenergic activity with propranolol administration increased the probability of win-stay compared to vehicle control (**Figure 2H**, GLMM, main effect of drug, p<0.0001, ꞵ1 = −0.023). There was also a significant interaction between sex and drug that males showed a greater increase in win-stay on propranolol compared to females (**Figure 2H inset,** GLMM, interaction term, 0.02, ꞵ3=-0.036). Interestingly, propranolol administration also decreased the probability of lose-shift but only in males (**Figure 2I**, GLMM, main effect of drug, 0.45, ꞵ1 = −0.01, interaction term, p= 0.002, ꞵ3 =0.08). Outcome sensitivity analysis revealed a significant increase in outcome sensitivity with propranolol administration (**Figure 1-1C**, GLMM, main effect of drug, 0.003, ꞵ1 = −0.186). Together, these results suggested an opposing role of dopamine and norepinephrine in modulating reward-driven decision making. Increasing dopamine or decreasing norepinephrine activity resulted in increased win-stay and decreased lose-shift but this effect is more sex-different in the noradrenergic manipulations.

### A Hidden Markov model (HMM) identifies distinct patterns of exploration and exploitation associated with modulation of dopamine and norepinephrine activity

The changes in the reward-driven behaviors such as win-stay and lose-shift could be a manifestation of strategic changes in how animals explored the changing environment. To understand how modulation of dopamine and norepinephrine affect the balance between exploration and exploitation, we used a Hidden Markov model (HMM) that modeled exploration and exploitation as two latent goal states to infer which choices were exploratory or exploitative (**Figure 3A)**. The HMM model has previously been used in rodents, primates and humans (Ebitz et al., 2018, 2020; Chen et al., 2021b) to infer explore-exploit state from choices and has shown robust correlation with neural activity as well as other behavioral metrics, including response time, value function, and reinforcement learning. We have previously established fast-switching (putatively explore) and slow-switching (putatively exploit) regimes in mice choice behavior (Chen et al., 2021b). This HMM method produces explore/exploit labels that better track neural activity and choice behavior across species than reinforcement learning (RL) models (Ebitz et al., 2018; Chen et al., 2021b). Figure 3A showed an example of HMM labeling of choices of an animal in the restless bandit task. The shaded area indicates exploratory trials inferred by the HMM. The animal displayed mixtures of exploratory bouts where choices were distributed across two choices and exploitative bouts where one choice was repeated.

**Figure 3.**
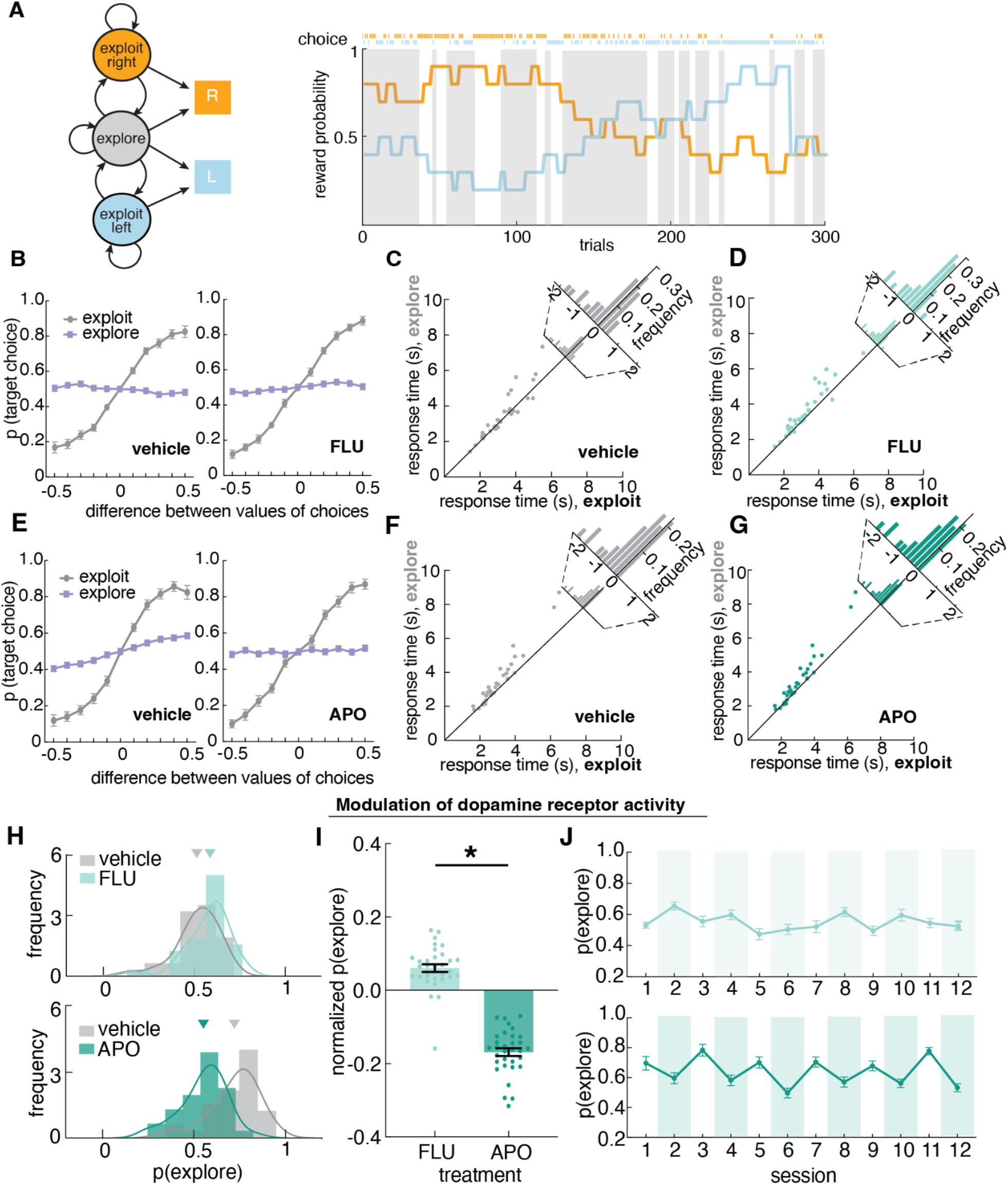
Up- and down-regulation of dopamine activity had bidirectional effects on the level of exploration. **A)** (left) Structure of a Hidden Markov model (HMM) that modeled exploration and exploitation as latent goal states underlying observed choices. This model incorporates two exploit states for left and right choice respectively, and one explore state where choices were uniformly distributed across options. (right) Reward probabilities, choices, and HMM labels for an example session of 300 trials for an example mouse. Shaded areas demonstrated HMM-labeled exploratory choices. **B)** Probability of choice as a function of value differences between choices for exploratory and exploitative states under vehicle (left) and flupenthixol (FLU) (right). **C, D)** Difference in response time for explore and exploit choices under vehicle and flupenthixol (FLU). **E)** Probability of choice as a function of value differences between choices for exploratory and exploitative states under vehicle (left) and apomorphine (APO) (right). **F, G)** Difference in response time for explore and exploit choices under vehicle and apomorphine (APO). **H)** Distribution of the percentage of HMM-labeled exploratory choices under flupenthixol (FLU)/ vehicle (top) and apomorphine (APO)/vehicle (bottom). **I)** Probability of exploration for dopamine antagonist (FLU) and agonist (APO), normalized by their vehicle control. Decreasing dopamine activity increased exploration and increasing dopamine activity decreased exploration. **J)** Probability of exploration by session with vehicle and drug session interleaved (flupenthixol: top; apomorphine: bottom). Drug administration sessions are in colored shades. * indicates p < 0.05. Graphs depict mean ± SEM across animals.

The HMM allows us to make statistical inferences about the probability that each choice was due to exploration or exploitation by modeling explore/exploit as the distinct latent goal states underlying choices. To evaluate the face validity of the HMM labels, we next examined whether HMM-labeled exploration matched the normative definition of exploration. First, by definition, exploration is a pattern of non-reward-maximizing choices whose purpose is to gain information. A non-reward maximizing goal would produce choices that are orthogonal to reward value. Therefore, we examined the probability of choosing a choice with regards to its relative value. In all control and dopamine treatment conditions, HMM-labeled exploratory choices were non-reward-maximizing: they were orthogonal to reward value, whereas exploit choices were correlated with relative reward value (**Figure 3B, E)**. The probability of choosing a target choice across bins of reward value was not different from chance (t(43) = 0.39, p = 0.70), which means that choice behavior was not driven by reward value during exploration. During exploration, animals chose both high-value choices and low-value choices at around chance level (mean = 49.8% ± 3.4% STD across all value bins). During exploitation, animals were much more likely to choose a high value choice over a low value choice (mean probability of choosing high value choice = 75.8% ± 10.0% STD, mean probability of choosing low value choice = 23.3% ± 10.1% STD). Second, prior research has shown that exploratory choices are more computationally expensive thus take longer than exploitative choices (Kool et al., 2010; Filipowicz et al., 2019; Chen et al., 2021b). We calculated the response time across HMM-inferred states and demonstrated that across all treatment conditions, HMM-inferred exploratory choices had longer response time than exploit choices (**Figure 3C, D, F, G,** paired t test, vehicle control (apomorphine): t(31) = 4.48, p<0.0001; apomorphine: t(31)= 4.42, p<0.0001; vehicle control (flupenthixol): t(31)=2.27, p=0.03; flupenthixol: t(31)=4.49, p<0.0001). Together, these results suggested that HMM produced meaningful and robust labeling of explore/exploit choices that match the normative definition of exploration and were able to explain variances in value selection and response time. Because this approach to infer exploration is agnostic to the generative models and depends only on the temporal statistics of choices (Ebitz et al., 2018, 2020; Chen et al., 2021b), it is particularly ideal for circumstances like this one, where we suspect that the generative computations may differ across treatment groups.

### Modulating dopamine receptor activity bidirectionally affected the level of exploration

To examine how much animals were exploring, we calculated the average number of exploratory choices inferred from HMM. The results revealed a bidirectional modulatory effect of dopamine receptor activity on the level of exploration. When animals were administered apomorphine, we found that animals on average had fewer exploratory trials than when they were on vehicle control, with 55.5% ± 12.6% STD of trials labeled as exploratory on apomorphine and 72.3% ± 13.6% STD of trials labeled as exploratory on vehicle (**Figure 3H, J,** GLMM, main effect of drug, p<0.0001, ꞵ1 = 0.174). This is also consistent with our findings in the win-stay lose-shift analysis that apomorphine increased the stickiness or repetitiveness of choice behaviors. In contrast, when decreasing dopamine receptor activity with flupenthixol administration, animals explore more compared to vehicle control, with 58.2% ± 12.0% STD of trials being exploratory on flupenthixol and 52.1% ± 12.3% STD of trials being exploratory on vehicle control (**Figure 3H, J**, GLMM, main effect of drug, 0.002, ꞵ1 = −0.062). This result suggested that manipulating dopamine receptor activity bidirectionally affected the level of exploration across both sexes (**Figure 3I**, **Figure 3-1)** - increasing dopamine receptor activity decreased exploratory choices and decreasing dopamine receptor activity increased exploratory choices.

### The effect of beta-noradrenergic receptor activity on exploration was sex-modulated

First, we conducted an HMM model validation by examining the value selection and response time during HMM-labeled explore and exploit state. Same as our findings in the dopamine treatment conditions, we found that in all norepinephrine conditions, HMM-inferred exploratory choices were also orthogonal to reward and exploit choices proportional to relative reward value (**Figure 4A, D)**. The probability of choosing a target choice across bins of reward value was not different from chance (t(43) = 0.85, p = 0.398). During exploration, animals chose both high-value choices and low-value choices at around chance level (mean = 49.8% ± 1.8% STD across all value bins). During exploitation, animals were much more likely to choose a high value choice over a low value choice (mean probability of choosing high value choice = 74.5% ± 8.6% STD, mean probability of choosing low value choice = 24.2% ± 9.3% STD). Response time during HMM-labeled exploratory choices were also longer than exploitative choices across all control conditions and propranolol condition (**Figure 4B, C, E, F,** paired t test, vehicle control (isoproterenol): t(31)=3.01, p=0.005; isoproterenol: t(31)=1.48, p=0.15; vehicle control (propranolol): t(31)=1.93, p=0.06; propranolol: t(30)=5.70, p<0.0001). With confidence in the approach to infer exploration, we next examined the effect of propranolol and isoproterenol on the level of exploration.

**Figure 4.**
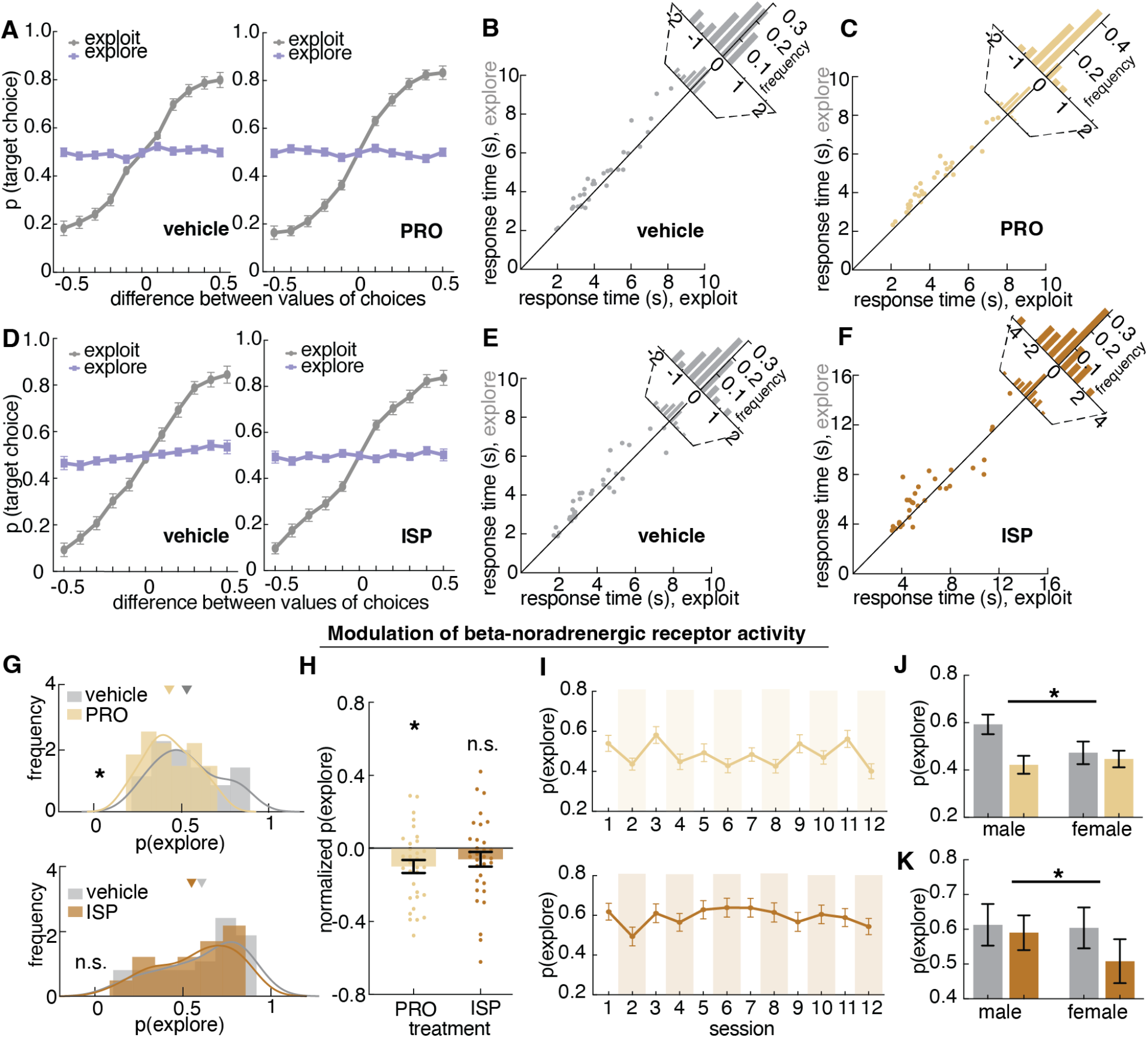
Downregulating norepinephrine activity influenced exploration but the effect was modulated by sex. **A)** Probability of choice as a function of value differences between choices for exploratory and exploitative states under vehicle (left) and propranolol (PRO) (right). **B, C)** Difference in response time for explore and exploit choices under vehicle and propranolol (PRO). **D)** Probability of choice as a function of value differences between choices for exploratory and exploitative states under vehicle (left) and isoproterenol (ISP) (right). **E, F)** Difference in response time for explore and exploit choices under vehicle and isoproterenol (ISP). **F)** Distribution of the percentage of HMM-labeled exploratory choices under propranolol (PRO)/ vehicle (top) and under isoproterenol (ISP)/vehicle (bottom). **H)** Probability of exploration for beta-noradrenergic antagonist (PRO) and agonist (ISP), normalized by their vehicle control. Propranolol decreased the level of exploration. **I)** Probability of exploration by session with vehicle and drug session interleaved. Drug administration sessions are in colored shades. Top: propranolol (PRO) condition; bottom: isoproterenol (ISP) condition. **J)** Probability of exploration under PRO and vehicle across sexes. The effect of PRO on exploration was primarily driven by a significant decrease in exploration under PRO in males. **K)** Probability of exploration under ISP and vehicle across sexes. ISP significantly decreased exploration in females but not males. * indicates p < 0.05. Graphs depict mean ± SEM across animals.

Modulating beta-noradrenergic receptor activity also influenced the probability of exploration and the modulatory effect on exploration was associated with sex. Decreasing beta-noradrenergic receptor activity with propranolol significantly decreased the level of exploration (**Figure 4G-I**) but only in males (**Figure 4J**, **Figure 3-1**, GLMM, interaction term, 0.0001, ꞵ1 = 0.138). Increasing beta-noradrenergic receptor activity with isoproterenol administration significantly decreased exploration in females (**Figure 3K**, **Figure 3-1**, interaction term, p= 0.002, ꞵ3=-0.103). Together these results suggested a sex-linked neuromodulatory effect of beta-noradrenergic receptor on exploration - increasing beta-noradrenergic activity decreased exploration in males and decreasing beta-noradrenergic activity decreased exploration in females. One possibility is that this sex-differentiated modulatory effect reflected sex-dependent ceiling/floor effects of noradrenergic signaling.

### Reinforcement learning (RL) models revealed changes in distinct parameters under modulation of dopamine and epinephrine

The results of the HMM suggested that both dopamine and norepinephrine modulation influenced the level of exploration. However, it is unclear whether changes in exploration were due to changes in similar or distinct latent cognitive parameters. Does dopamine and norepinephrine modulate exploration via distinct or overlapping mechanisms? To address this question, we fitted a series of reinforcement learning (RL) models to understand the effect of pharmacological manipulation on the latent cognitive parameters that could influence exploration and exploitation (Ishii et al., 2002; Daw et al., 2006; Pearson et al., 2009; Jepma and Nieuwenhuis, 2011).

To make inferences about changes in the RL model parameters, we first identified the best-fitting RL model for the animals’ behaviors. The majority of RL model parameters can be categorized in three ways: value-dependent learning terms, value-independent bias terms, and decision noise/randomness terms (Katahira, 2018). Previous studies demonstrated the effect of various RL parameters on value-based decision making, including value-dependent terms such as learning rate and asymmetrical learning rate (Frank et al., 2007; Gershman, 2016; Cinotti et al., 2019; Chen et al., 2021b), value-independent terms such as choice bias (Katahira, 2018; Wilson and Collins, 2019; Chen et al., 2021b), and noise terms such as inverse temperature and lapse rate (Economides et al., 2015; Cinotti et al., 2019; Wilson and Collins, 2019). Here, we compared six reinforcement learning models that incorporated one or more of the above parameters (learning rate, bias, noise) and had different assumptions about the latent cognitive processes that animals might employ during exploration (**Figure 5A**, **Equations 3-7)**. These models included: (1) a “random” model with some overall bias for one choice over the other, (2) a “noisy win-stay lose-shift” model (WSLS) that assumes a win-stay lose-shift policy with some level of randomness, (3) a two-parameter “RL” model with a consistent learning rate and some inverse temperature that captures decision noise, (4) a three-parameter “RLCK” model that captures both value-based and value-independent decision with two parameters for learning rate and choice bias, and an overall decision noise parameter, (5) a three-parameter “RLγ” model that captures asymmetrical learning with a learning rate parameter and a scaling parameter for negative outcome learning and a decision noise parameter, and (6) a four-parameter “RLCKγ” asymmetrical learning and bias model that includes a choice bias term on top of the “RLγ” (see methods). We fitted these models to each individual animal and compared the likelihood of each of the six models under four vehicle control conditions and four drug conditions.

**Figure 5.**
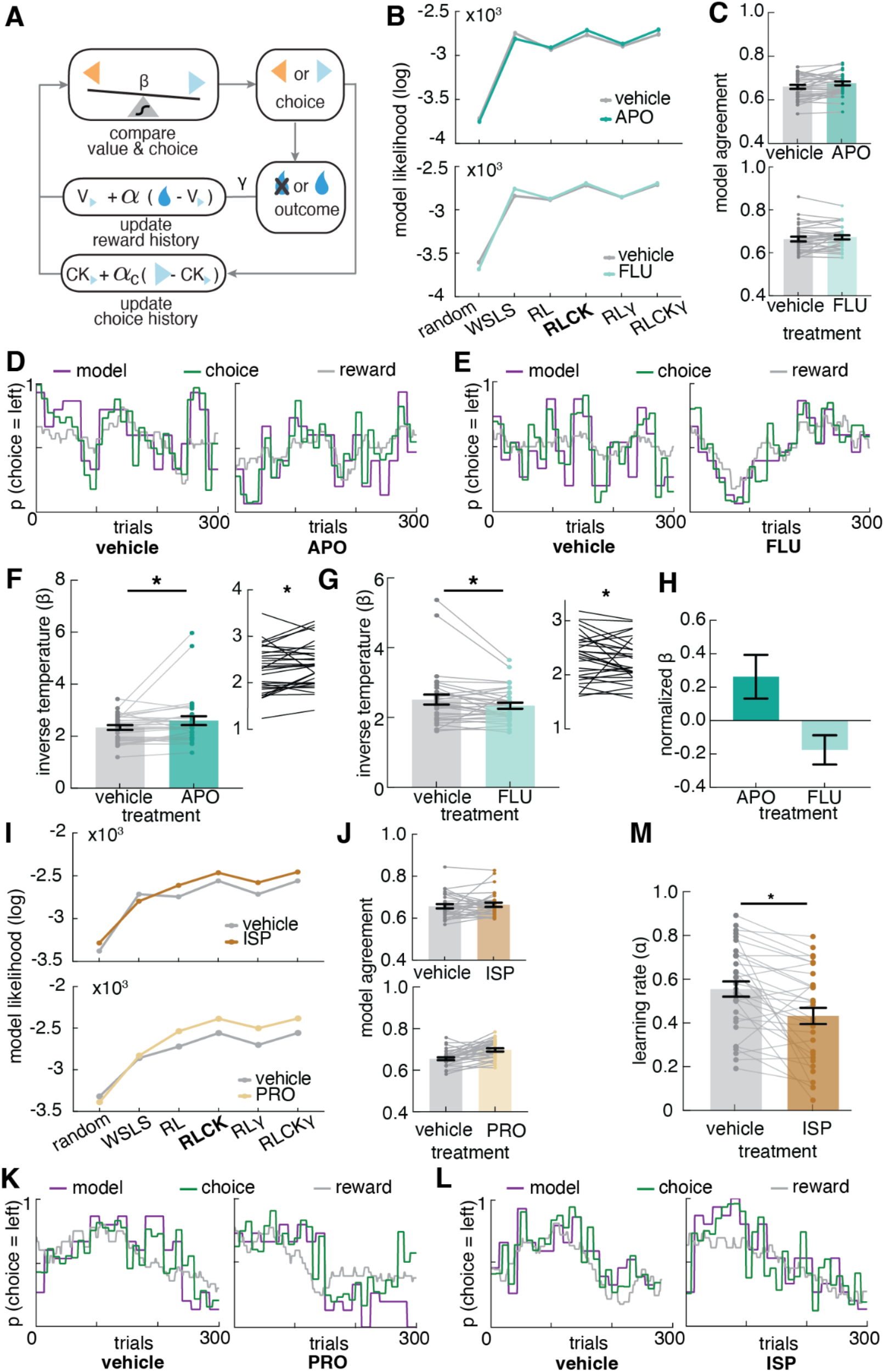
Modulation of dopamine and noradrenergic receptor activity led to changes in different reinforcement learning (RL) model parameters. **A)** A diagram of reinforcement learning (RL) parameters that capture learning rate (α), asymmetric learning (γ), choice bias (α_c_), inverse temperature/decision noise (β). The RL models tested included one or more of the above parameters. **B)** Model likelihood for six RL models using log likelihood for vehicle and apomorphine (APO) condition (top), and vehicle and flupenthixol condition (bottom). The three-parameter RLCK model was the best fit and most parsimonious model for behaviors. **C)** Model agreement for the best fit model (RLCK), which measures the probability of the model predicting the actual choices of animals for vehicle and apomorphine (APO) condition (top), and vehicle and flupenthixol condition (bottom). **D, E)** Actual choices (purple) and simulated choices (green) from the best fit model (RLCK) for the same animal under vehicle/apomorphine and vehicle/flupenthixol. Grey line represents the reward probability of the left choice. **F)** Average inverse temperature (β) across apomorphine and vehicle. (inset) inverse temperature across conditions after removing two outliers. **G)** Average inverse temperature (β) across flupenthixol and vehicle. (inset) inverse temperature across conditions after removing two outliers. **H**) Agonizing (apomorphine) and antagonizing (flupenthixol) dopamine activity revealed bidirectional effect on inverse temperature, controled by their own vehicle control. **I**) Model likelihood for six RL models using log likelihood for vehicle and isoproterenol (ISP) condition (top) and vehicle and propranolol (PRO) condition (bottom). **J)** Model agreement for the best fit model (RLCK), which measures the probability of the model predicting the actual choices of animals for vehicle and isoproterenol (ISP) condition (top) and vehicle and propranolol (PRO) condition (bottom). **K, L)** Actual choices (purple) and simulated choices (green) from the best fit model (RLCK) for the same animal under vehicle/propranolol and vehicle/isoproterenol. Grey line represents the reward probability of the left choice. **M)** Average learning rate (α) across isoproterenol and vehicle. * indicates p < 0.05. Graphs depict mean ± SEM across animals.

In dopamine modulation conditions, the RLCK model (learning + choice kernel,) and the RLCKγ (asymmetrical learning and bias) model best fit the behaviors (**Figure 5B**, average AIC across DA conditions: model 1 random: 73838.03; model 2 WSLS: 55843.26; model 3 RL: 58133.45; model 4 RLCK: 54652.02; model 5 RLγ: 57545.47; model 6 RLCKγ: 54612.56). Since the RLCKγ model did not significantly improve the model fitting, we decided to use the more parsimonious RLCK model that has three parameters (learning rate, choice bias, and decision noise). To examine how well the best fit RLCK model was at actually predicting animals’ choices, we measured the model agreement for each model, which was calculated as the probability of choice correctly predicted by the optimized model parameters for each model (**Figure 5C**). In dopamine agonist condition, the RLCK model predicted over 66% of animals’ actual choices (vehicle: 66.1% ± 5.3% STD, APO: 67.6% ± 5.0%). In the dopamine antagonist condition, the RLCK model also predicted over 66% of animals’ actual choices (vehicle: 66.4% ± 6.4%, FLU: 67.3% ± 5.4%). Figure 5D and 5E showed the model simulation, actual choice, and reward probability of the same animal. This also visually demonstrated that the RLCK model was able to accurately predict animal’s changing choice behaviors. Additionally, this also demonstrated that animal choice behaviors were following the reward contingency.

Then we examined how the reinforcement learning model parameters changed with dopamine modulation. Previous study has shown that both learning rate and inverse temperature parameters could induce changes in the overall level of exploration (Chen et al., 2021b). Therefore, we asked whether the bidirectional changes in exploration when up- or down-regulating dopamine receptor activity were due to changes in learning rate or decision noise. The results revealed a bidirectional effect of dopamine modulation on the inverse temperature parameter (**Figure 5F-H**). Because RL model data can be non-normally distributed, the Shapiro-Wilk test of normality was conducted to determine whether the RL parameters were normally distributed. The result suggested that the parameters were not always normally distributed (decision noise: vehicle: p= 0.85, APO: p<0.0001; vehicle: p<0.0001, FLU: p=0.09), therefore we will report both parametric and non-parametric statistics results for RL parameters. Increasing dopamine receptor activity with apomorphine administration increased inverse temperature and decreased random noise (**Figure 5F**, Wilcoxon matched-pairs test, p= 0.0165; paired t-test, p= 0.053). Decreasing dopamine receptor activity with flupenthixol administration decreased inverse temperature and increased random noise (**Figure 5G**, Wilcoxon matched-pairs test, p= 0.0253; paired t-test, p= 0.053). Neither dopamine agonist (apomorphine) nor dopamine antagonist (flupenthixol) significantly influenced learning rate or choice kernel compared to vehicle controls.

In norepinephrine modulation conditions, the RLCK model (learning + choice kernel,) and the RLCKγ (asymmetrical learning and bias) model also best fit the behavioral data (**Figure 5I**, average AIC across NE conditions: model 1 random: 66898.83; model 2 WSLS: 56061.3; model 3 RL: 53194.59; model 4 RLCK: 50035.02; model 5 RLγ: 52664.48; model 6 RLCKγ:50027.82). However, since the RLCKγ model did not significantly improve the model fitting, we decided to use the more parsimonious RLCK model for the NE condition as well. The model agreement was calculated for all NE modulations. In the noradrenergic agonist condition, the RLCK model predicted over 65% of animals’ actual choices (**Figure 5J**, vehicle: 65.7% ± 5.9% STD, ISP: 66.4% ± 5.7%). In the noradrenergic antagonist condition, the RLCK model also predicted over 65% of animals’ actual choices (vehicle: 65.5% ± 3.9% STD, PRO: 69.8% ± 4.5%). Figure 5K and 5L showed the model simulation, actual choice, and reward probability of the same animal as Figure 5D and 5E.

We found that modulating beta-noradrenergic receptor activity also resulted in changes in the reinforcement learning model parameter. Up-regulating norepinephrine activity with isoproterenol significantly decreased the learning rate (**Figure 5M**, Wilcoxon matched-pairs test, p= 0.0001; paired t-test, p= 0.0008). This result is consistent with the decreased win-stay and decreased lose-shift behaviors when administered isoproterenol because the lower learning rate under isoproterenol resulted in reduced outcome sensitivity (**Supplementary figure 1C**), which means learning less from either a rewarded or a non-rewarded outcome. There were no significant changes in other parameters when administered isoproterenol compared to vehicle control. We did not find any significant changes in any reinforcement learning parameters when administered propranolol compared to vehicle control. Together, these results suggested distinct roles of dopaminergic drugs in modulating exploration via decision noise and isoproterenol in modulating exploration via outcome sensitivity.

## Discussion

In this study, we pharmacologically up- and down-regulated dopamine and beta-noradrenergic receptor activity and examined the modulatory effects of the manipulations on exploration in an explore/exploit task. We used a combination of computational modeling approaches to characterize changes in the level of exploration and latent cognitive variables that could contribute to exploration. Modulating dopamine receptor activity revealed a bi-directional modulatory effect on exploration - increasing dopamine receptor activity decreased the level of exploration and decreasing dopamine receptor activity increased the level of exploration.

Modulation of beta-noradrenergic receptor activity also influenced how much animals explored - beta-noradrenergic receptor antagonist (propranolol) decreased the level of exploration. However, when examining the modulatory effect of beta-noradrenergic receptor activity on exploration across sexes, we found that modulatory effect was sex-different: isoproterenol selectively decreased exploration in females whereas propranolol decreased exploration in males. Previously, the role of dopamine and norepinephrine in mediating exploration were often examined in isolation with different task designs. This study allowed the direct comparison of the modulatory effect of dopamine and norepinephrine on the transition between exploration and exploitation in the same task across the same animals.

Dopamine’s function as a neuromodulator has been described in two key ways. First, the phasic activation of midbrain dopamine neurons is thought to be a key contributor to reinforcement learning (Barto, 1995; Montague et al., 1995, 1996; Schultz et al., 1997; Schultz, 1998). Numerous studies across species have shown that dopamine neurons encoded the reward prediction error (RPE) (Schultz, 1998; Niv, 2009). Ventral tegmental dopamine neuronal activity and release in ventral striatum is proposed to reflect this error, which is used to update action values (Schultz, 1998, 2013; Niv, 2009). Consistent with this view, some previous pharmacological investigations in human decision making have observed changes in model-derived learning rates that directly correlate with dopamine synthesis agonism via L-DOPA (Frank et al., 2004; Rutledge et al., 2009) and dopamine receptor D2/D3 antagonism via amisulpride (Cremer et al., 2022). However, this framework does not necessarily account for a large body of literature describing a second way to view dopamine’s neuromodulatory function, as tonic levels influencing motivation, vigor, and cognitive flexibility (Floresco, 2013; Beeler et al., 2014), especially when pharmacologically targeted in the frontal cortex. Because pharmacological agents are by definition modulating receptor activity over long periods of time, this role for tonic dopamine is poised to be computationally described. Indeed, when pharmacologically or genetically modulating tonic dopamine function, many studies have noted profound changes in the decision “noise” (inverse temperature) or perseverative errors - consistent with a role for dopamine modulation in cognitive flexibility in earlier work (Beeler et al., 2010; Humphries et al., 2012; Eisenegger et al., 2014; Lee et al., 2015; Cinotti et al., 2019; Ebitz et al., 2019). Two prior examples combining pharmacological manipulation and computational modeling in animal tasks explicitly testing the explore-exploit tradeoff particularly highlight this. Cinotti et al. found that systemic dopamine blockade with flupenxithol increased the amount of random exploratory choices without affecting learning rate in a rat three-arm bandit task (Cinotti et al., 2019), particularly in situations where uncertainty was manipulated. Similarly, Ebitz et al. reported that systemic cocaine, which blocks dopamine reuptake, reduced flexibility and regulated tonic exploration in rhesus macaques (Ebitz et al., 2019). Likewise, we find that systemic activation of dopamine receptors significantly decreased exploration as defined by our Hidden Markov model and decision noise as described by our reinforcement learning model, while antagonism of these receptors increased these computational parameters measuring flexible behavior. Our data thus provide additional support for the idea that dopamine modulation across the brain provides more than a learning signal, but also a key signal tuning the precision or flexibility of behavior.

As noted in our introduction, the roles of dopamine and norepinephrine are sometimes conflated in the computational neuroscience literature. This is in part because modulation of these systems can be correlated, and noninvasive measures of catecholamine function, for example blink rate and pupil diameter, could be ascribed to the endogenous activity of either neuromodulator (although pharmacological action may target one or the other) (Jepma and Nieuwenhuis, 2011; Ebitz and Moore, 2018; Sescousse et al., 2018; Findling et al., 2019; Cremer et al., 2022). This is further complicated by the fact that some studies examining the relationship between dopamine and exploration have used pharmacological manipulations which simultaneously influence both tonic dopamine and norepinephrine levels, such as psychostimulants or L-DOPA (Pessiglione et al., 2006; Rutledge et al., 2009; Orsini et al., 2016; Ebitz et al., 2019; Chakroun et al., 2020). This makes it challenging to dissociate the modulatory roles of dopamine and norepinephrine in exploration i.e. how dopamine and norepinephrine distinctively contribute to changes in reinforcement learning model parameters, and addressing this challenge was a major goal of our study.

Surprisingly, we found that when modulating beta-noradrenergic receptor activity changed the level of exploration, it did so via changes in outcome sensitivity. Previous studies have proposed the locus coeruleus-norepinephrine (LC-NE) system to regulate exploration by changing decision noise. Kane et al. chemogenetically stimulated locus coeruleus activity in rats and found that increased LC tonic activity increased in increased exploration, best explained by an increase in decision noise (Kane et al., 2017; Wilson et al., 2021). Other studies have echoed the findings that norepinephrine levels predicted the level of random exploration, or decision noise, in the system (Jepma and Nieuwenhuis, 2011; Ebitz and Moore, 2018; Joshi and Gold, 2020). However, in this study we did not find any changes in the noise parameter (inverse temperature, or how closely you adhere to your value rule) associated with up- and down-regulation of beta-noradrenergic activity. Instead, the results revealed some learning rate changes when increasing beta-noradrenergic receptor activity. Model-free analysis on outcome sensitivity suggested that increasing or decreasing beta-noradrenergic receptor activity bi-directionally changed outcome sensitivity. Propranolol increased outcome sensitivity and decreased exploration, which means the animals were more likely to switch choice after loss and more likely to repeat rewarded choices. In contrast isoproterenol decreased outcome sensitivity, as seen in reduced win-stay and lose-shift behavior. This is consistent with the decreased learning rate when administered isoproterenol because a decrease in learning rate is associated with decreased value learning from each outcome. Where might this strange association between noradrenergic modulation, exploration, and outcome sensitivity emerge from? We have previously shown that exploration as defined by our Hidden Markov model can be driven by either changes in learning rate or changes in inverse temperature (Chen et al., 2021b), supporting the idea that the exploration we see here could be driven by long-term changes in the influence of outcomes. Work focused on beta noradrenergic signaling in the context of aversive learning has highlighted a role for noradrenergic signaling in extinction (Iordanova et al., 2021), over a phasic timeframe of minutes to hours. This suggests the intriguing possibility that the effect of noradrenergic blockade here may be to prevent extinction of learned values. In short, although the most common associations in the computational neuroscience literature are between dopamine and reward learning versus norepinephrine and decision noise, our current data suggests a more nuanced role for each of these modulators.

Adding an additional layer of nuance to these findings, we find that sex differences are a major axis of variability in the effect of beta-noradrenergic receptor modulation on the explore-exploit tradeoff. Increasing NE receptor activity with the beta agonist isoproterenol decreased exploration only in males, while decreasing NE receptor activity with the beta antagonist propranolol decreased exploration especially in females. One possible explanation is that the relationship between exploration and NE receptor activity was nonlinear but instead, followed an inverted U-shape. Such inverted U-shape correlation has been suggested between cognitive performance and NE level (Aston-Jones and Cohen, 2005; Redish et al., 2007; Mäki-Marttunen et al., 2020). Since females have been reported to have higher tonic NE levels due to both function and structural differences (Pinos et al., 2001; Bangasser et al., 2016), it is possible that the baseline difference in tonic NE level across sexes could posit individuals on different parts of the inverted U-curve, where increasing or decreasing NE could affect exploration in one sex without impacting the other.

Taken together, this study demonstrates the distinct role of dopamine and norepinephrine in tuning the exploration-exploitation tradeoff when learning about an uncertain environment. Dopamine modulated exploration via decision noise and norepinephrine modulated exploration via outcome sensitivity. The systemic manipulation of dopamine and norepinephrine receptor activity did not allow for more precise modulation of phasic changes in catecholamine levels. However, the computational tool that we presented here can complement higher resolution neural recording techniques to examine for phasic changes in dopamine and norepinephrine activity on a trial-by-trial basis. Finally, this study also highlighted that the modulatory effects of catecholamine on exploration-exploitation tradeoff are strongly influenced by sex and sex-linked mechanisms. Therefore, future research should take into account sex-correlated individual variability when examining the neural correlates of exploration.

## Acknowledgements

This work was supported by NIMH R01 MH123661 (N.M.G), NIMH P50 MH119569 (N.M.G), and NIMH T32 training grant MH115886 (C.S.C), startup funds from the University of Minnesota (N.M.G.), a Young Investigator Grant from the Brain and Behavior Foundation (R.B.E.), an Unfettered Research Grant from the Mistletoe Foundation (R.B.E.), and Fonds de Recherche du Québec Santé, Chercheur-Boursier Junior 1, #284309 (R.B.E). Thank you to Dr. David Redish and Madison J Merfeld for comments improving this manuscript. Thank you to Anila Bano for helping with the experiment.

---

**Supplementary figure 1.**
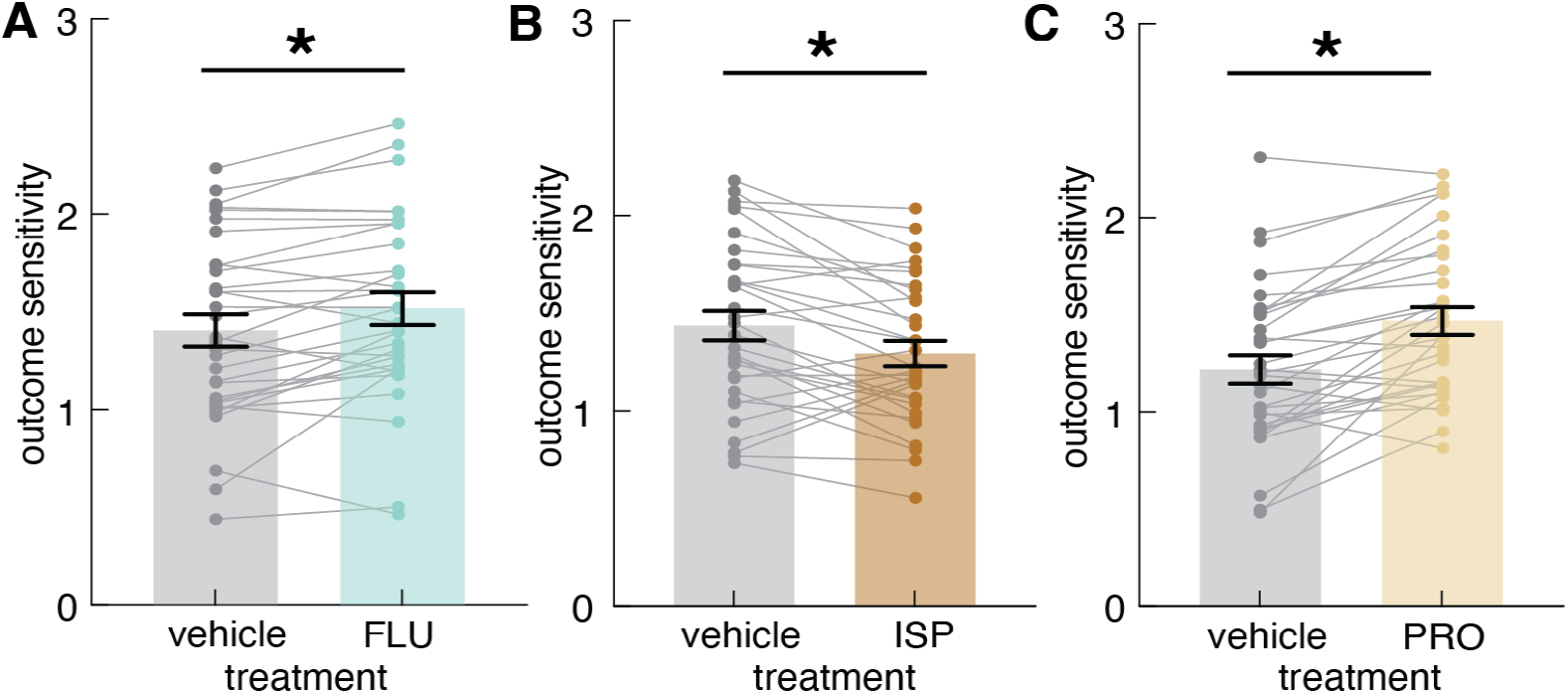
Figure 1-1. Effect of dopamine and norepinephrine modulation on outcome sensitivity. Related to Figure 1. **A)** Average outcome sensitivity across flupenxithol (FLU) and vehicle. **B**) Average outcome sensitivity across isoproterenol (ISP) and vehicle. **C**) Average outcome sensitivity across propranolol (PRO) and vehicle. * indicates p < 0.05. Graphs depict mean ± SEM across animals.

**Supplementary figure 2.**
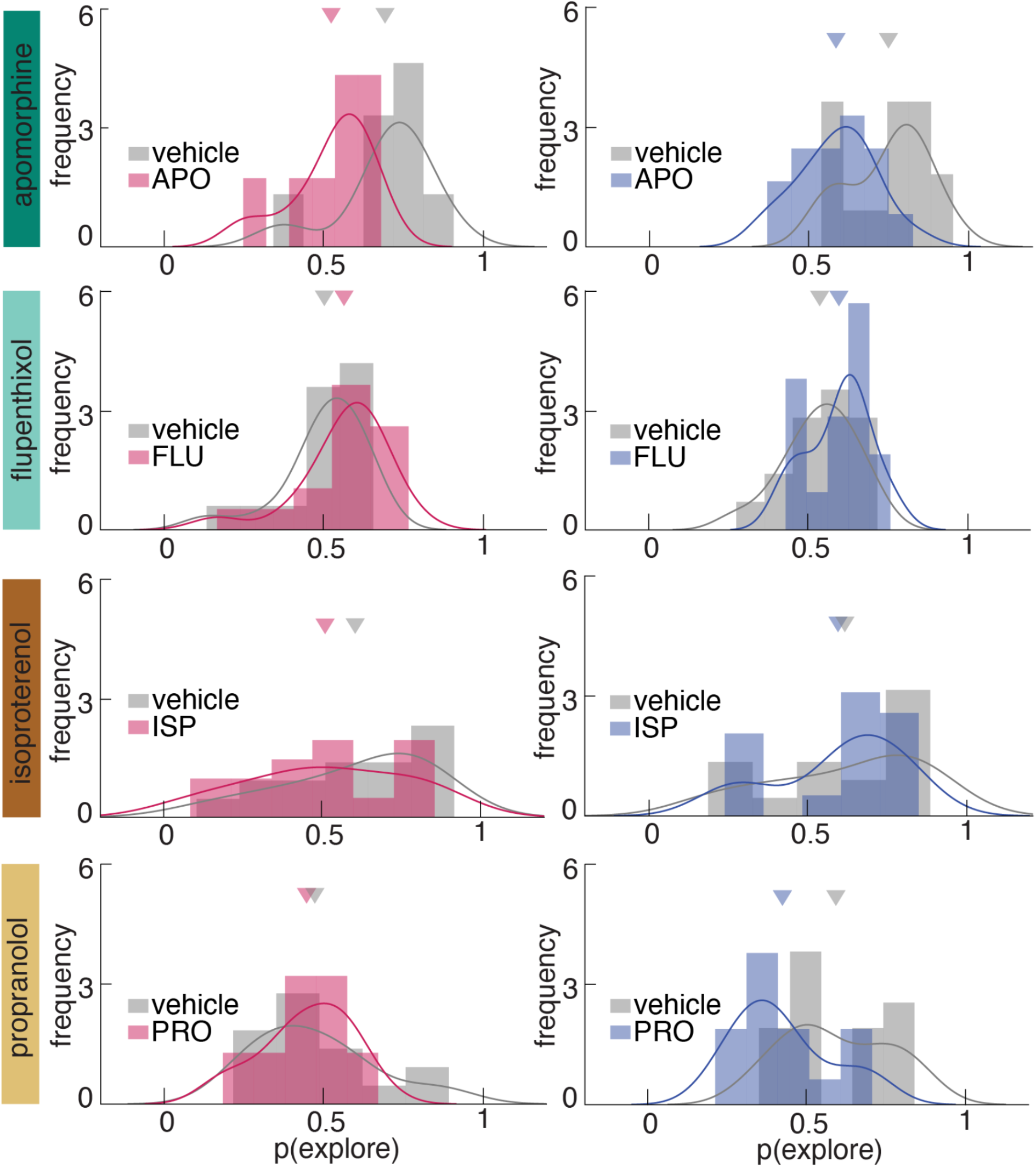
Figure 3-1. Effect of dopamine and norepinephrine modulation on exploration in female and male mice. Related to Figure 3,4.

## References

Addicott MA, Pearson JM, Schechter JC, Sapyta JJ, Weiss MD, Kollins SH (2020) Attention-deficit/hyperactivity disorder and the explore/exploit trade-off. Neuropsychopharmacology Available at: http://dx.doi.org/10.1038/s41386-020-00881-8.

Amemiya S, Kubota N, Umeyama N, Nishijima T, Kita I (2016) Noradrenergic signaling in the medial prefrontal cortex and amygdala differentially regulates vicarious trial-and-error in a spatial decision-making task. Behav Brain Res 297:104–111 Available at: http://dx.doi.org/10.1016/j.bbr.2015.09.002.

Aston-Jones G, Cohen JD (2005) An integrative theory of locus coeruleus-norepinephrine function: adaptive gain and optimal performance. Annu Rev Neurosci 28:403–450 Available at: http://dx.doi.org/10.1146/annurev.neuro.28.061604.135709.

Bangasser DA, Wiersielis KR, Khantsis S (2016) Sex differences in the locus coeruleus-norepinephrine system and its regulation by stress. Brain Res 1641:177–188 Available at: http://dx.doi.org/10.1016/j.brainres.2015.11.021.

Barto AG (1995) Adaptive critics and the basal ganglia. Models of information processing in the basal ganglia 382:215–232 Available at: https://psycnet.apa.org/fulltext/1995-97527-007.pdf.

Beeler JA, Cools R, Luciana M, Ostlund SB, Petzinger G (2014) A kinder, gentler dopamine… highlighting dopamine’s role in behavioral flexibility. Front Neurosci 8:4 Available at: http://dx.doi.org/10.3389/fnins.2014.00004.

Beeler JA, Daw N, Frazier CRM, Zhuang X (2010) Tonic dopamine modulates exploitation of reward learning. Front Behav Neurosci 4:170 Available at: http://dx.doi.org/10.3389/fnbeh.2010.00170.

Bilmes J (1998) A Gentle Tutorial of the EM Algorithm and its Application to Parameter Estimation for Gaussian Mixture and Hidden Markov Models. Available at: http://citeseerx.ist.psu.edu/viewdoc/summary?doi=10.1.1.28.613 [Accessed April 23, 2021].

Bouret S, Sara SJ (2005) Network reset: a simplified overarching theory of locus coeruleus noradrenaline function. Trends Neurosci 28:574–582 Available at: http://dx.doi.org/10.1016/j.tins.2005.09.002.

Cabib S, Puglisi-Allegra S, Oliverio A (1984) Chronic stress enhances apomorphine-induced stereotyped behavior in mice: involvement of endogenous opioids. Brain Res 298:138–140 Available at: http://dx.doi.org/10.1016/0006-8993(84)91156-9.

Chakroun K, Mathar D, Wiehler A, Ganzer F, Peters J (2020) Dopaminergic modulation of the exploration/exploitation trade-off in human decision-making. Elife 9 Available at: http://dx.doi.org/10.7554/eLife.51260.

Chen CS, Ebitz RB, Bindas SR, Redish AD, Hayden BY, Grissom NM (2021a) Divergent Strategies for Learning in Males and Females. Curr Biol 31:39–50.e4 Available at: http://dx.doi.org/10.1016/j.cub.2020.09.075.

Chen CS, Knep E, Han A, Ebitz RB, Grissom N (2021b) Sex differences in learning from exploration. Elife 10 Available at: http://dx.doi.org/10.7554/eLife.69748.

Cinotti F, Fresno V, Aklil N, Coutureau E, Girard B, Marchand AR, Khamassi M (2019) Dopamine blockade impairs the exploration-exploitation trade-off in rats. Sci Rep 9:6770 Available at: http://dx.doi.org/10.1038/s41598-019-43245-z.

Cohen JD, McClure SM, Yu AJ (2007) Should I stay or should I go? How the human brain manages the trade-off between exploitation and exploration. Philosophical Transactions of the Royal Society B: Biological Sciences 362:933–942 Available at: http://dx.doi.org/10.1098/rstb.2007.2098.

Cremer A, Kalbe F, Müller JC, Wiedemann K, Schwabe L (2022) Disentangling the roles of dopamine and noradrenaline in the exploration-exploitation tradeoff during human decision-making. Neuropsychopharmacology Available at: http://dx.doi.org/10.1038/s41386-022-01517-9.

Daw ND, O’Doherty JP, Dayan P, Seymour B, Dolan RJ (2006) Cortical substrates for exploratory decisions in humans. Nature 441:876–879 Available at: http://dx.doi.org/10.1038/nature04766.

Ebitz RB, Albarran E, Moore T (2018) Exploration Disrupts Choice-Predictive Signals and Alters Dynamics in Prefrontal Cortex. Neuron 97:475 Available at: http://dx.doi.org/10.1016/j.neuron.2018.01.011.

Ebitz RB, Moore T (2018) Both a Gauge and a Filter: Cognitive Modulations of Pupil Size. Front Neurol 9:1190 Available at: http://dx.doi.org/10.3389/fneur.2018.01190.

Ebitz RB, Sleezer BJ, Jedema HP, Bradberry CW, Hayden BY (2019) Tonic exploration governs both flexibility and lapses. PLoS Comput Biol 15:e1007475 Available at: http://dx.doi.org/10.1371/journal.pcbi.1007475.

Ebitz RB, Tu JC, Hayden BY (2020) Rules warp feature encoding in decision-making circuits. PLoS Biol 18:e3000951 Available at: http://dx.doi.org/10.1371/journal.pbio.3000951.

Economides M, Kurth-Nelson Z, Lübbert A, Guitart-Masip M, Dolan RJ (2015) Model-Based Reasoning in Humans Becomes Automatic with Training. PLoS Comput Biol 11:e1004463 Available at: http://dx.doi.org/10.1371/journal.pcbi.1004463.

Eisenegger C, Naef M, Linssen A, Clark L, Gandamaneni PK, Müller U, Robbins TW (2014) Role of dopamine D2 receptors in human reinforcement learning. Neuropsychopharmacology 39:2366–2375 Available at: http://dx.doi.org/10.1038/npp.2014.84.

Elliott R (2003) Executive functions and their disorders. Br Med Bull 65:49–59 Available at: http://dx.doi.org/10.1093/bmb/65.1.49.

Fernandez-Tome MP, Sanchez-Blazquez P, del Rio J (1979) Impairment by apomorphine of one-trial passive avoidance learning in mice: the opposing roles of the dopamine and noradrenaline systems. Psychopharmacology 61:43–47 Available at: http://dx.doi.org/10.1007/BF00426809.

Filipowicz ALS, Levine J, Piasini E, Tavoni G, Kable JW, Gold JI (2019) The complexity of model-free and model-based learning strategies. Available at: http://dx.doi.org/10.1101/2019.12.28.879965.

Findling C, Skvortsova V, Dromnelle R, Palminteri S, Wyart V (2019) Computational noise in reward-guided learning drives behavioral variability in volatile environments. Nat Neurosci 22:2066–2077 Available at: http://dx.doi.org/10.1038/s41593-019-0518-9.

Floresco SB (2013) Prefrontal dopamine and behavioral flexibility: shifting from an “inverted-U” toward a family of functions. Front Neurosci 7:62 Available at: http://dx.doi.org/10.3389/fnins.2013.00062.

Frank MJ, Moustafa AA, Haughey HM, Curran T, Hutchison KE (2007) Genetic triple dissociation reveals multiple roles for dopamine in reinforcement learning. Proc Natl Acad Sci U S A 104:16311–16316 Available at: http://dx.doi.org/10.1073/pnas.0706111104.

Frank MJ, Seeberger LC, O’reilly RC (2004) By carrot or by stick: cognitive reinforcement learning in parkinsonism. Science 306:1940–1943 Available at: http://dx.doi.org/10.1126/science.1102941.

Gershman SJ (2016) Empirical priors for reinforcement learning models. J Math Psychol 71:1–6 Available at: https://www.sciencedirect.com/science/article/pii/S0022249616000080.

Gershman SJ (2019) Uncertainty and Exploration. Decision (Wash D C) 6:277–286 Available at: http://dx.doi.org/10.1037/dec0000101.

Goldschmidt PL, Frances H, Simon P (1984) Stimulation of beta-adrenergic receptors and spontaneous motor activity in mice. Pharmacol Biochem Behav 21:177–180 Available at: http://dx.doi.org/10.1016/0091-3057(84)90210-7.

Grigoryan GA (2012) Serotonin and Impulsivity (animal experiments). Neurosci Behav Physiol 42:885–894 Available at: https://doi.org/10.1007/s11055-012-9654-3.

Grissom NM, Reyes TM (2019) Let’s call the whole thing off: evaluating gender and sex differences in executive function. Neuropsychopharmacology Available at: https://www.nature.com/articles/s41386-018-0179-5.

Grossman CD, Bari BA, Cohen JY (2020) Serotonin neurons modulate learning rate through uncertainty. bioRxiv:2020.10.24.353508 Available at: https://www.biorxiv.org/content/10.1101/2020.10.24.353508v1 [Accessed September 29, 2021].

Humphries MD, Khamassi M, Gurney K (2012) Dopaminergic Control of the Exploration-Exploitation Trade-Off via the Basal Ganglia. Front Neurosci 6:9 Available at: http://dx.doi.org/10.3389/fnins.2012.00009.

Ichihara K, Nabeshima T, Kameyama T (1988) Opposite effects induced by low and high doses of apomorphine on single-trial passive avoidance learning in mice. Pharmacol Biochem Behav 30:107–113 Available at: http://dx.doi.org/10.1016/0091-3057(88)90431-5.

Iordanova MD, Yau JO-Y, McDannald MA, Corbit LH (2021) Neural substrates of appetitive and aversive prediction error. Neurosci Biobehav Rev 123:337–351 Available at: http://dx.doi.org/10.1016/j.neubiorev.2020.10.029.

Ishii S, Yoshida W, Yoshimoto J (2002) Control of exploitation-exploration meta-parameter in reinforcement learning. Neural Netw 15:665–687 Available at: http://dx.doi.org/10.1016/s0893-6080(02)00056-4.

Jepma M, Nieuwenhuis S (2011) Pupil Diameter Predicts Changes in the Exploration– Exploitation Trade-off: Evidence for the Adaptive Gain Theory. J Cogn Neurosci 23:1587– 1596 Available at: https://doi.org/10.1162/jocn.2010.21548.

Jepma M, Schaaf JV, Visser I, Huizenga HM (2020) Uncertainty-driven regulation of learning and exploration in adolescents: A computational account. PLoS Comput Biol 16:e1008276 Available at: http://dx.doi.org/10.1371/journal.pcbi.1008276.

Joshi S, Gold JI (2020) Pupil Size as a Window on Neural Substrates of Cognition. Trends Cogn Sci 24:466–480 Available at: http://dx.doi.org/10.1016/j.tics.2020.03.005.

Kane GA, Vazey EM, Wilson RC, Shenhav A, Daw ND, Aston-Jones G, Cohen JD (2017) Increased locus coeruleus tonic activity causes disengagement from a patch-foraging task. Cogn Affect Behav Neurosci 17:1073–1083 Available at: http://dx.doi.org/10.3758/s13415-017-0531-y.

Kaske EA, Chen CS, Meyer C, Yang F, Ebitz B, Grissom N, Kapoor A, Darrow DP, Herman AB (2022) Prolonged physiological stress is associated with a lower rate of exploratory learning that is compounded by depression. Biol Psychiatry Cogn Neurosci Neuroimaging Available at: https://linkinghub.elsevier.com/retrieve/pii/S2451902222003391.

Katahira K (2018) The statistical structures of reinforcement learning with asymmetric value updates. J Math Psychol 87:31–45 Available at: https://www.sciencedirect.com/science/article/pii/S0022249617302407.

Kim H, Lee D, Shin Y-M, Chey J (2007) Impaired strategic decision making in schizophrenia. Brain Res 1180:90–100 Available at: http://dx.doi.org/10.1016/j.brainres.2007.08.049.

Kobayashi K (2001) Role of catecholamine signaling in brain and nervous system functions: new insights from mouse molecular genetic study. J Investig Dermatol Symp Proc 6:115–121 Available at: http://dx.doi.org/10.1046/j.0022-202x.2001.00011.x.

Kool W, McGuire JT, Rosen ZB, Botvinick MM (2010) Decision making and the avoidance of cognitive demand. J Exp Psychol Gen 139:665–682 Available at: http://dx.doi.org/10.1037/a0020198.

Lee E, Seo M, Dal Monte O, Averbeck BB (2015) Injection of a dopamine type 2 receptor antagonist into the dorsal striatum disrupts choices driven by previous outcomes, but not perceptual inference. J Neurosci 35:6298–6306 Available at: http://dx.doi.org/10.1523/JNEUROSCI.4561-14.2015.

Mäki-Marttunen V, Andreassen OA, Espeseth T (2020) The role of norepinephrine in the pathophysiology of schizophrenia. Neurosci Biobehav Rev 118:298–314 Available at: http://dx.doi.org/10.1016/j.neubiorev.2020.07.038.

Malloy-Diniz LF, Miranda DM, Grassi-Oliveira R (2017) Editorial: Executive Functions in Psychiatric Disorders. Front Psychol 8:1461 Available at: http://dx.doi.org/10.3389/fpsyg.2017.01461.

Money KM, Stanwood GD (2013) Developmental origins of brain disorders: roles for dopamine. Front Cell Neurosci 7:260 Available at: http://dx.doi.org/10.3389/fncel.2013.00260.

Montague PR, Dayan P, Person C, Sejnowski TJ (1995) Bee foraging in uncertain environments using predictive hebbian learning. Nature 377:725–728 Available at: http://dx.doi.org/10.1038/377725a0.

Montague PR, Dayan P, Sejnowski TJ (1996) A framework for mesencephalic dopamine systems based on predictive Hebbian learning. J Neurosci 16:1936–1947 Available at: https://www.ncbi.nlm.nih.gov/pubmed/8774460.

Morris LS, Baek K, Kundu P, Harrison NA, Frank MJ, Voon V (2016) Biases in the Explore-Exploit Tradeoff in Addictions: The Role of Avoidance of Uncertainty. Neuropsychopharmacology 41:940–948 Available at: http://dx.doi.org/10.1038/npp.2015.208.

Mussey JL, Travers BG, Klinger LG, Klinger MR (2015) Decision-making skills in ASD: performance on the Iowa Gambling Task. Autism Res 8:105–114 Available at: http://dx.doi.org/10.1002/aur.1429.

Nakamura K, Kurasawa M, Tanaka Y (1998) Apomorphine-induced hypoattention in rats and reversal of the choice performance impairment by aniracetam. Eur J Pharmacol 342:127– 138 Available at: http://dx.doi.org/10.1016/s0014-2999(97)01457-x.

Niv Y (2009) Reinforcement learning in the brain. J Math Psychol 53:139–154 Available at: https://www.princeton.edu/~yael/Publications/Niv2009.pdf.

Niv Y, Joel D, Meilijson I, Ruppin E (2002) Evolution of reinforcement learning in uncertain environments: A simple explanation for complex foraging behaviors. Adapt Behav 10:5–24 Available at: https://psycnet.apa.org/fulltext/2003-02884-002.pdf.

Orsini CA, Willis ML, Gilbert RJ, Bizon JL, Setlow B (2016) Sex differences in a rat model of risky decision making. Behav Neurosci 130:50–61 Available at: http://dx.doi.org/10.1037/bne0000111.

Pearson JM, Hayden BY, Raghavachari S, Platt ML (2009) Neurons in posterior cingulate cortex signal exploratory decisions in a dynamic multioption choice task. Curr Biol 19:1532–1537 Available at: http://dx.doi.org/10.1016/j.cub.2009.07.048.

Pessiglione M, Seymour B, Flandin G, Dolan RJ, Frith CD (2006) Dopamine-dependent prediction errors underpin reward-seeking behaviour in humans. Nature 442:1042–1045 Available at: http://dx.doi.org/10.1038/nature05051.

Pinos H, Collado P, Rodríguez-Zafra M, Rodríguez C, Segovia S, Guillamón A (2001) The development of sex differences in the locus coeruleus of the rat. Brain Res Bull 56:73–78 Available at: http://dx.doi.org/10.1016/s0361-9230(01)00540-8.

Ranjbar-Slamloo Y, Fazlali Z (2019) Dopamine and Noradrenaline in the Brain; Overlapping or Dissociate Functions? Front Mol Neurosci 12:334 Available at: http://dx.doi.org/10.3389/fnmol.2019.00334.

Redish AD, Jensen S, Johnson A, Kurth-Nelson Z (2007) Reconciling reinforcement learning models with behavioral extinction and renewal: implications for addiction, relapse, and problem gambling. Psychol Rev 114:784–805 Available at: http://dx.doi.org/10.1037/0033-295X.114.3.784.

Ressler KJ, Nemeroff CB (2001) Role of norepinephrine in the pathophysiology of neuropsychiatric disorders. CNS Spectr 6:663–666, 670 Available at: http://dx.doi.org/10.1017/s1092852900001358.

Rothenhoefer KM, Costa VD, Bartolo R, Vicario-Feliciano R, Murray EA, Averbeck BB (2017) Effects of Ventral Striatum Lesions on Stimulus-Based versus Action-Based Reinforcement Learning. J Neurosci 37:6902–6914 Available at: http://dx.doi.org/10.1523/JNEUROSCI.0631-17.2017.

Rutledge RB, Lazzaro SC, Lau B, Myers CE, Gluck MA, Glimcher PW (2009) Dopaminergic drugs modulate learning rates and perseveration in Parkinson’s patients in a dynamic foraging task. J Neurosci 29:15104–15114 Available at: http://dx.doi.org/10.1523/JNEUROSCI.3524-09.2009.

Schultz W (1998) Predictive reward signal of dopamine neurons. J Neurophysiol 80:1–27 Available at: http://dx.doi.org/10.1152/jn.1998.80.1.1.

Schultz W (2013) Updating dopamine reward signals. Curr Opin Neurobiol 23:229–238 Available at: http://dx.doi.org/10.1016/j.conb.2012.11.012.

Schultz W, Dayan P, Montague PR (1997) A neural substrate of prediction and reward. Science 275:1593–1599 Available at: http://dx.doi.org/10.1126/science.275.5306.1593.

Seamans JK, Yang CR (2004) The principal features and mechanisms of dopamine modulation in the prefrontal cortex. Prog Neurobiol 74:1–58 Available at: http://dx.doi.org/10.1016/j.pneurobio.2004.05.006.

Sescousse G, Ligneul R, van Holst RJ, Janssen LK, de Boer F, Janssen M, Berry AS, Jagust WJ, Cools R (2018) Spontaneous eye blink rate and dopamine synthesis capacity: preliminary evidence for an absence of positive correlation. Eur J Neurosci 47:1081–1086 Available at: http://dx.doi.org/10.1111/ejn.13895.

Sutton RS, Barto AG (1998) Reinformcent Learning: An Introduction.

Warren CM, Wilson RC, van der Wee NJ, Giltay EJ, van Noorden MS, Cohen JD, Nieuwenhuis S (2017) The effect of atomoxetine on random and directed exploration in humans. PLoS One 12:e0176034 Available at: http://dx.doi.org/10.1371/journal.pone.0176034.

Weed MR, Gold LH (1998) The effects of dopaminergic agents on reaction time in rhesus monkeys. Psychopharmacology 137:33–42 Available at: http://dx.doi.org/10.1007/s002130050590.

Williams OOF, Coppolino M, George SR, Perreault ML (2021) Sex Differences in Dopamine Receptors and Relevance to Neuropsychiatric Disorders. Brain Sci 11 Available at: http://dx.doi.org/10.3390/brainsci11091199.

Wilson RC, Bonawitz E, Costa VD, Ebitz RB (2021) Balancing exploration and exploitation with information and randomization. Curr Opin Behav Sci 38:49–56 Available at: http://dx.doi.org/10.1016/j.cobeha.2020.10.001.

Wilson R, Collins A (2019) Ten simple rules for the computational modeling of behavioral data. Available at: https://pdfs.semanticscholar.org/91b9/d3ab7532ea24ae70cd726355f25235b1fe8b.pdf.

Yu AJ, Dayan P (2005) Uncertainty, neuromodulation, and attention. Neuron 46:681–692 Available at: http://dx.doi.org/10.1016/j.neuron.2005.04.026.

